# Tuning non-linear mechanics in collagen hydrogels modulates cellular morphotypes in three dimensions

**DOI:** 10.1101/2024.03.18.585457

**Authors:** Marco A. Enriquez Martinez, Zhao Wang, Robert J. Ju, Petri Turunen, Jitendra Mata, Elliot P. Gilbert, Jan Lauko, Samantha J. Stehbens, Alan E. Rowan

**Affiliations:** Australian Institute for Bioengineering and Nanotechnology (AIBN), The University of Queensland, Brisbane QLD 4072, Australia; Institute for Molecular Biosciences (IMB), The University of Queensland, Brisbane QLD 4072, Australia; Institute of Molecular Biology, Mainz, Germany; Australian Centre for Neutron Scattering, Australian Nuclear Science and Technology Organisation, Lucas Height, NSW 2234, Australia

## Abstract

Collagen networks contribute to tissue architecture and modulate cellular responses in crowded three-dimensional environments. Therefore, it is the most widely used biological polymer in three-dimensional studies of cellular interactions with the extracellular matrix. *In vivo*, collagen exists embedded within additional matrix components. Studies have shown that the combination of matrices induces synergistic mechanical interactions, influencing the non-linear mechanical behaviour of collagen networks. However, how cells respond to changes in collagen non-linear elasticity remains largely unknown. By precisely controlling the mechanical behaviour of collagen networks with the biologically inert and semiflexible polymer polyisocyanopeptides, we demonstrate that changes in the non-linear elasticity of collagen induces morphological cell responses that influence how cells migrate, proliferate, and interact with collagen. We found that when collagen rigidifies in the presence of a second component, this induces morphological changes in cell-matrix interactions, resulting in a decrease in migration and the ability of cells to deform collagen matrices. Our results demonstrate that the onset of collagen stiffening is key to inducing intracellular tension which dictates morphological cell responses in three-dimensional collagen networks. We anticipate our findings will prove useful in understanding how cells respond to changes in collagen mechanics when combined in double network systems which better recapitulates tissues *in vivo*.

## Introduction

The architecture of mammalian tissues is maintained by the constant secretion and degradation of collagen.^1^ From a mechanical perspective, this constant production of collagen provides tissues such as skin, lungs and cartilage with resistance against mechanical stretching and compression by stiffening.^2–5^ The nature of this mechanical behaviour arises from the non-linear elasticity of collagen networks.^6^ At the tissue level, an extensive body of evidence recognises that the constant stiffening of collagenous tissues is an important part of tissue growth. Mechanical loading by exercise stimulates collagen synthesis,^7^ while increased tension within collagen fibrils ensures resistance against enzymatic degradation.^8,9^ Nevertheless, stiffening of collagen also arises from within the interstitial matrix by cellular movement and remodelling. Cells stiffen collagen to generate traction in mesenchymal-based migration^10,11^ and to communicate over long distances.^12–14^

During disease or injury, such as cancer progression and wound healing, new extracellular matrix is excessively deposited into the interstitial space at a faster rate than which it is degraded.^15–17^ This mechanism is important for the survival and growth of cancer cells to drive metastasis,^18^ and to facilitate the rapid generation of scar tissue during wound closure.^19^ In these scenarios, it is often reported that tissues rigidify due to an increase in the deposition of collagen and intrafibrillar crosslinking between collagen fibrils.^15,20,21^ However, other alterations in the interstitial matrix that also lead to the rigidification of tissues include the large deposition of fibrin^22^ and hyaluronic acid.^23,24^ Mechanical models of composite systems demonstrate that the presence of these two components influences the ability of collagen fibres to bend and stiffen under load, requiring higher levels of stress for collagen networks to stiffen.^25–30^ However, an unanswered question within the field of biological soft matter is how these changes in collagen’s non-linear elasticity can modulate cellular behaviour.

In this study, we investigated biological responses of cancer cells and fibroblasts to changes in collagen’s non-linear elasticity by developing a collagen composite system with polyisocyanopeptides (PIC).^31^ By precisely tuning the non-linear mechanical behaviour of collagen hydrogels with low PIC concentrations and controllable contour lengths, we identified that the parameter that governs the response of cells to the extracellular environment in three-dimensions is the onset of collagen’s stiffening (σ_*c*_). Our results show that, as collagen slowly rigidifies and loses sensitivity towards stress, cells exhibit morphological and behavioural changes consistent with an increase in cellular tension. We demonstrate that cells exposed to rigid collagen networks have a reduced ability to form cellular extensions correlating with a decrease in cell migration persistence and a loss of the active deformation of the surrounding matrix. We further found that these biological responses are concomitant with a biphasic response of integrin-based adhesions to mechanical load. In that, as collagen rigidifies, cells at first productively engage with cell-matrix adhesions that grow and elongate at the lateral regions of the cell’s leading edge. However, when we shift the onset of collagen’s stiffening (σ_*c*_) to high levels of stress beyond a threshold, cells fail to deform the collagen matrix. As a result, the interaction between cells and the collagen network weakens. This suggests that the onset of collagen stiffening (σ_*c*_) modulates the force loading rate within focal adhesions in 3D collagen networks. Here we present a novel collagen composite system capable of precisely tuning the non-linear elasticity of collagen. We provide new insights into the mechanical parameters that govern physical interactions between cells and collagen matrices. Application of our novel collagen composite system will generate a new understanding of how cells respond to the rigidification of collagen, giving insight to homeostatic tissue maintenance as well as pathological conditions, such as fibrosis and cancer.

## Results

### PIC tunes the gelation properties of collagen hydrogels with controllable polymer density and contour length

To control the mechanical behaviour of collagen hydrogels, we generated a series of collagen composites where we combined cold solutions of polyisocyanopeptides (PIC)^31^ with cold solutions of atelo type I collagen before the reconstitution process (Fig. 1a). In this composite system, the collagen concentration was kept constant (2 mg mL^−1^, pH 7, Phosphate Buffered Saline, 1X PBS), while the network parameters of PIC were tuned by increasing the PIC density (0.2–2.0 mg mL^−1^) or contour length (*L_c_*) of the polymer (low molecular weight; LMW, S, and high molecular weight; HMW, L).^32,33^

**Fig 1.**
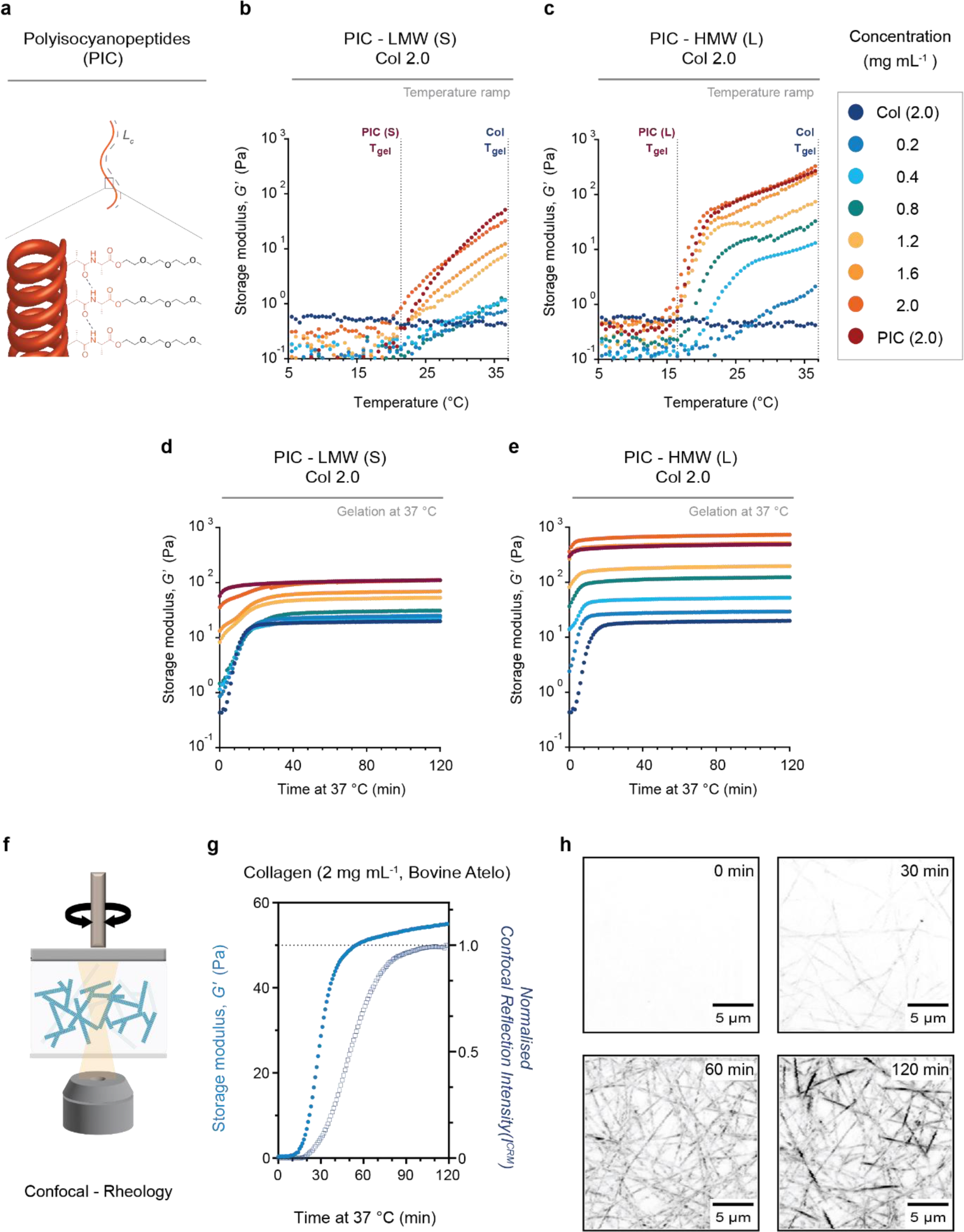
Mechanical properties of PIC Collagen composites. **a,** schematic representation of polyisocyanopeptides (PIC). Contour length *L_c_* is shown in gray dotted lines as the end-to-end distance along the polymer backbone. **b,c**, linear temperature ramps from 5 to 37 °C (1 °C min^−1^) showing the influence of PIC on the onset of gelation temperature (*T_gel_*) of collagen type I solutions (bovine atelo, 1X PBS, pH 7) of constant concentration (2 mg mL^−1^) with solutions of increasing PIC concentrations (0.2 – 2.0 mg mL^−1^, 1X PBS) and different PIC molecular weight (Low Molecular Weight, LMW, M_v_: 283 kg mol^−1^; and High Molecular Weight, HMW, M_v_: 518 kg mol^−1^). **b**, LMW (S) PIC and **c**, HMW (L) PIC. Gel point (T_gel_) is defined as the onset in the storage modulus (*G’*, Pa). Both gelation profiles show a shift in T_gel_ below physiological temperature and a rapid increase in *G’*. **d,e**, gelation profiles of composite hydrogels incubated *in situ* between plates for 2 hours at 37 °C after temperature ramps. All sets of data are an average of three independent measurements. **f**, schematic of a combined laser scan confocal microscope with a rheometer. Schematic of a collagen network is shown in blue. **g**, graph shows the evolution of the *sol-gel* transition (increase in storage modulus (*G’*) (Pa)) at 37 °C and the intensity profile in confocal reflection mode (*I^CRM^)* (normalised) of the pure bovine atelo collagen type I hydrogel (2 mg mL ^−1^, 1X PBS (Phosphate Buffered Saline), pH 7). Data is acquired by simultaneously applying a small oscillatory shear and acquiring time lapse images every 1 min. Graph represents data from an average of three independent measurements. **h**, inverted confocal images show the formation of the collagen network every 30 min until the *I^CRM^* profile reaches a plateau at ∼120 min. Z projections show 20 µm depth. Scale bar (5 µm). The increase in *I^CRM^* is correlated to the continuing thickening of collagen fibres by lateral growth until the hydrogel undergoes complete gelation.

To determine the influence of PIC on the mechanical behaviour of collagen hydrogels, we first investigated the effect of PIC on collagen’s gelation with oscillatory shear rheology experiments using a temperature ramp (5–37 °C; at 1 °C min^−1^) (Fig. 1 b,c). Our results show that even a minimal addition of the synthetic polymer (0.2–0.8 mg mL^−1^) is sufficient to influence the gelation temperature (*T_gel_*) of the collagen-based composites. Consistent with the mechanical profile of PIC,^32^ the *T_gel_* of the composites is dependent on PIC’s contour length. Collagen composites with LMW PIC (Fig. 1b) gelate at higher temperatures compared to composites with HMW PIC (Fig. 1c; ∼22 °C vs. ∼17 °C, respectively). In addition, increasing the density of PIC within a concentration range of 0.8–2.0 mg mL^−1^ results in a substantial increase in collagen’s linear moduli (storage modulus, *G’*) (100 –1,000 Pa). (Fig. 1 d,e; Supplementary Fig. 1). These rheology experiments demonstrate that PIC can tune the linear mechanical behaviour of collagen hydrogels without altering the temperature or collagen concentration.

### PIC and Collagen form an interpenetrating network (IPN) by self-assemblies driven by physiological heat

To ensure that the mechanical enhancement was not due to changes in collagen architecture, we next evaluated the formation of the collagen network in the presence of PIC. Having observed that the composites gelated at lower temperatures, we hypothesised that the formation of the collagen network was accelerated. Previous reports demonstrate that increasing the polymer density of a second network accelerates collagen fibrillogenesis via an effect of macromolecular crowding (MMC).^34^ ^35–38^ To test this, we first evaluated the formation of atelo type I collagen hydrogels by placing cold collagen solutions between a rheometer and an inverted laser scan microscope (Fig. 1 f-h). This allowed us to simultaneously monitor the formation of the collagen network structure with changes in its viscoelastic profile as the temperature was increased to 37 °C. Atelo collagen hydrogels undergo a phase transition from a solution to a viscoelastic gel (*sol*-*gel* transition) at 37 °C by showing an increase in storage moduli (*G’*, Pa) accompanied by a delayed increase in reflection intensity (intensity confocal reflectance mode, I*^CRM^*) (Fig. 1g), indicative of the slow formation of the network structure (Fig. 1h).

Next, we monitored the gelation and reflection intensity of collagen composites containing higher concentrations of PIC (Fig. 2 a-c). The lower refractive index of the PIC allowed us to solely monitor the formation of the collagen networks in the absence of any fluorescent labels by time-lapse imaging. In PIC–collagen composites, we observed an immediate increase in *G’* (Pa) at physiological temperature (37 °C) (Fig. 2b). The delayed increase in *I^CRM^*relative to *G’* (Pa) indicated that the reconstitution of the collagen network was not accelerated by the gelation of PIC, and that there were no changes in the duration of the reconstitution. Instead, the assembly of the collagen network was only initiated when exposed to physiological temperature (Fig. 2c; 0-120 min at 37 °C). Importantly, higher concentrations of PIC (0.8–2.0 mg mL^−1^) showed a similar trend (Supplementary Fig. 2) indicating that the increase in PIC density does not induce a macromolecular crowding effect.

**Fig 2.**
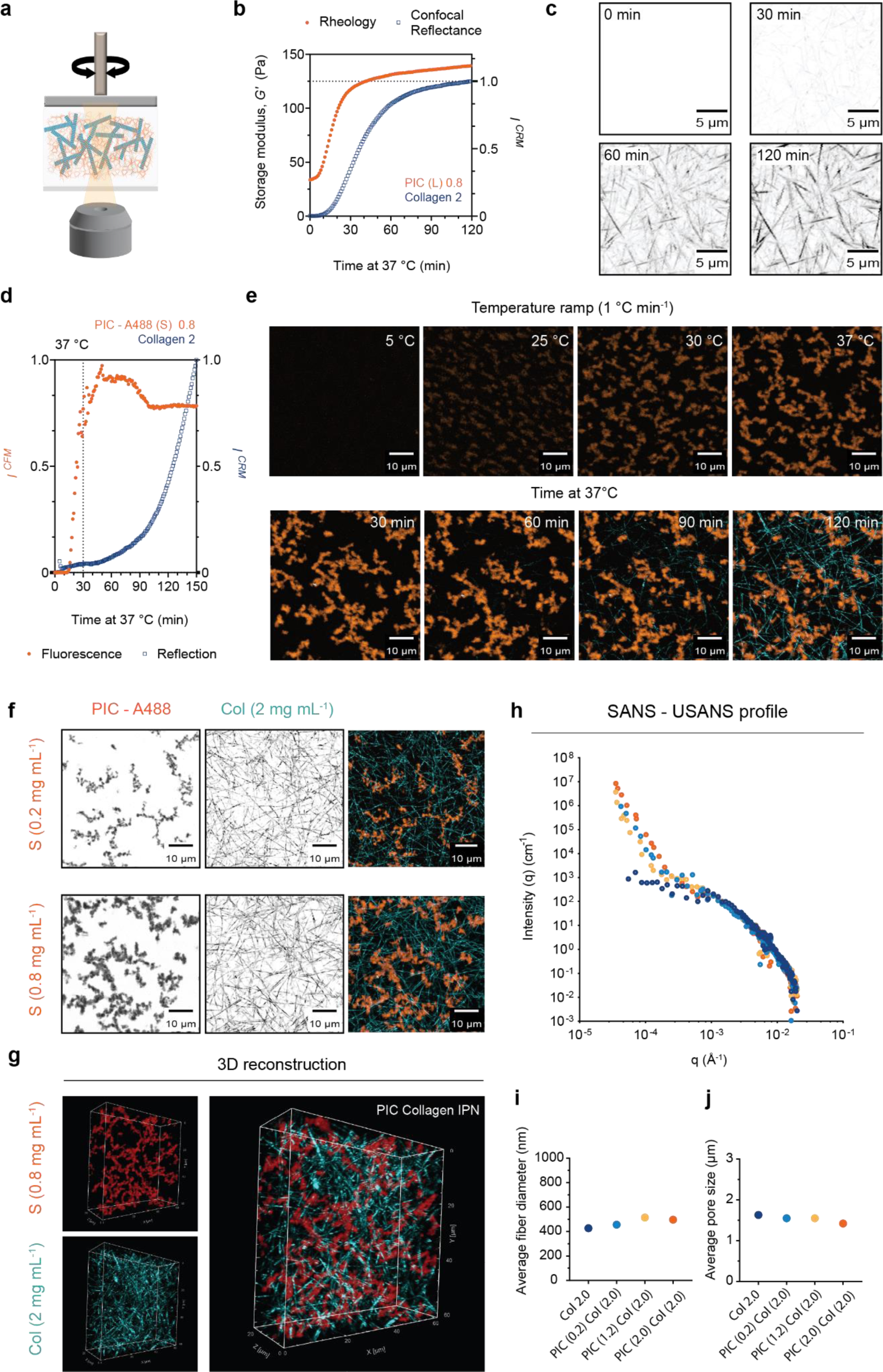
Elucidating collagen’s architecture in PIC Collagen interpenetrating networks. **a**, schematic of a combined confocal microscopy – rheology set up used to investigate the formation of the interpenetrating networks (IPN). Collagen network schematic is shown in blue and PIC network in orange. **b**, addition of PIC induces a rapid increase in *G’* (Pa) at 37 °C compared to single atelo collagen hydrogels. Graph shows the gelation of methoxy-functionalised PIC (L, 0.8 mg mL^−1^) with collagen (2 mg mL^−1^). Sets of data are representative from an average of two independent measurements. **c**, inverted confocal images show the formation of the atelo collagen network within the composite using confocal reflection mode. Z-projection shows 20 µm depth. Scale bar (5 µm). **d**, PIC-Collagen IPN follows a sequential order of IPN formation monitored by simultaneous reflectance and fluorescence microscopy. PIC (S, 1/100 azide) was fluorescently labelled with DBCO-Alexa 488 (PIC (S)-A488, 1/10 A488/Azide) and atelo collagen formation was monitored by reflection mode at λ = 647 nm. Graph shows fluorescent and reflection intensities (normalised) characteristic of each network as a function of temperature and time. Note that quantified reflection and fluorescence intensities are a representation from maximal projections of 10 µm depth. Graph shows data from an average of two independent measurements. **e**, gelation was induced with a linear temperature ramp from 5 to 37 °C (1 °C min^−1^) (Time 0 to 30 min) and temperature was kept constant at 37 °C for 2 hours (Time 30 to 150 min). Time lapse images (acquired every 1 min) show the formation of the PIC network from ∼ 25 °C to 37 °C and the subsequent formation of the collagen network after 120 min at 37 °C. Time lapse images show maximal projections of 10 µm depth. Scale bar (10 µm). **f,** maximal projections of PIC-Collagen IPN using different concentrations of PIC-A488 (S, 0.2 mg mL^−1^ and 0.8 mg mL^−1^). Confocal images show PIC-A488 (S, 0.2 and 0.8 mg mL^−1^) in fluorescence mode and atelo collagen (2.0 mg mL^−1^) in reflection mode. Images were acquired at 37 °C. Z projections show 10 µm depth. Scale bar (10 µm). **g**, three-dimensional (3D) reconstruction of a PIC–Collagen IPN (PIC (S) – A488; 0.8 mg mL^−1^ Col 2.0 mg mL^−1^) showing interpenetration between the networks. X-Y-Z stack shows 60 µm x 60 µm x 20 µm depth. **h,** combined Ultra Small Angle Neutron Scattering (USANS) – Small Angle Neutron Scattering (SANS) intensity profiles of atelo collagen (2.0 mg mL^−1^) and PIC–Collagen composites (S, 0.2, 1.2 2.0 mg mL^−1^). Samples were dissolved in 20% D_2_O (refer to methods for contrast variation experiments). Scattering data from Porod regions were fitted and analysed to determine the influence of PIC in collagen’s fibre diameter (**i**) and pore size (**j**). We observe no substantial difference in collagen’s pore size and fibre diameter at lower and higher q (Å^−1^) ranges in the presence of different PIC densities.

To visualise the order of network formation, we performed simultaneous time-lapse acquisitions by confocal fluorescence (CFM) and reflection (CRM) modes using confocal– rheology (Fig. 2 d,e). We labelled an azide-functionalised PIC (LMW) with Alexa Fluor® 488-DBCO (PIC-A488)^39^ and monitored the formation of both networks during the temperature ramp (5–37 °C). The fluorescence intensity (*I^CFM^*) emitted by PIC-A488 increased rapidly at lower temperatures showing a fast formation of the PIC network. The fluorescence intensity (*I^CFM^*) remained constant while the reflection intensity (*I^CRM^*) emitted by the collagen network increased only after being exposed to 37 °C (Fig. 2d). This indicates that the formation of the PIC network precedes the assembly of the collagen network.

To demonstrate that there was no phase separation between the materials once they were formed, we performed co-image analysis and three-dimensional reconstitution of the two networks (Fig. 2 f,g). We observed that the PIC was homogenously distributed and interpenetrated throughout the collagen network. To further quantitatively evaluate the architecture of the collagen network in the presence of PIC, we characterised the structural properties of collagen in the composites by small angle neutron scattering – ultra small angle scattering (SANS–USANS) (Fig. 2h) and confocal laser scan microscopy (CLSM) (Supplementary Fig. 3). Both methods demonstrate that the presence of PIC in the early stages of fibrillogenesis does not have an impact on collagen’s pore size or fibre diameter. In neutron scattering experiments, we first employed contrast variation experiments to eliminate the scattering signal emitted by PIC. We found the contrast match point of PIC at 20% D^2^O and determined that collagen’s diameter (400–500 nm) (Fig. 2i) and pore size (1.5 µm) (Fig. 2j) remained consistent compared to pure collagen hydrogels. Analysis of the collagen’s pore size by CLSM also showed a similar trend, with collagen’s pore size (1.5–2 µm) remaining unaffected by increasing the PIC concentration (Supplementary Fig. 3). Together, these results demonstrate that the mechanical properties of collagen hydrogels can be tuned by using a low weight percentage (wt%) of PIC without affecting collagen’s architecture.

### PIC controls the stress-stiffening response of collagen networks

After determining that the PIC does not influence collagen’s assembly or architecture, we next investigated how the semi flexibility of PIC can synergistically influence the response of atelo collagen networks to applied shear stress. To this end, we applied oscillatory shear stress *σ* (Pa) to the composites after gelation and evaluated changes in the non-linear differential modulus (*K’*), particularly the onset of stiffening (σ_*c*_) (Fig. 3 a-h). When networks of reconstituted collagen stiffen, they display a non-linear response to stress dominated by a bending (m ∼ 1) to stretching transition (m ∼ 1/ 2) in the non-linear regime.^6,40,41^ However, when combining collagen with LMW PIC, we first observed that, as the density of the LMW PIC increases, the onset of stiffening (*σ_c_*) of atelo collagen is shifted towards higher levels of stress (Fig. 3 c-e), meaning that atelo collagen becomes less responsive to stress. The shift in the non-linear response becomes only apparent when reaching concentrations of 0.8 and 1.2 mg mL^−1^ of LMW PIC (Fig. 3e). Coincidently, this range of PIC concentrations falls within the regime where PIC depicts a non-linear response dominated by entropic fibre stretching (m ∼ 3/2).^32^ Moreover, our experiments using HMW PIC show that one can also decrease the sensitivity of collagen to applied stress by increasing the rigidity of PIC bundles (Fig. 3 f-h). The same decrease in sensitivity observed in composites with high concentrations of LMW PIC (2 mg mL^−1^) can be achieved with lower concentrations of HMW PIC (0.8–1.2 mg mL^−1^). Re-scaling the stress-stiffening curves into a master curve indicates that all the composites are dominated by bending energy in the non-linear regime (*m* ∼ 1) (Supplementary Fig. 4 and 5), and that the entropic response (*m* ∼ 3/2) of PIC is suppressed even when adding high concentrations of HMW PIC. This suggests that PIC controls collagen’s stress-stiffening response by controlling the resistance of individual collagen fibres to bending.

**Fig 3.**
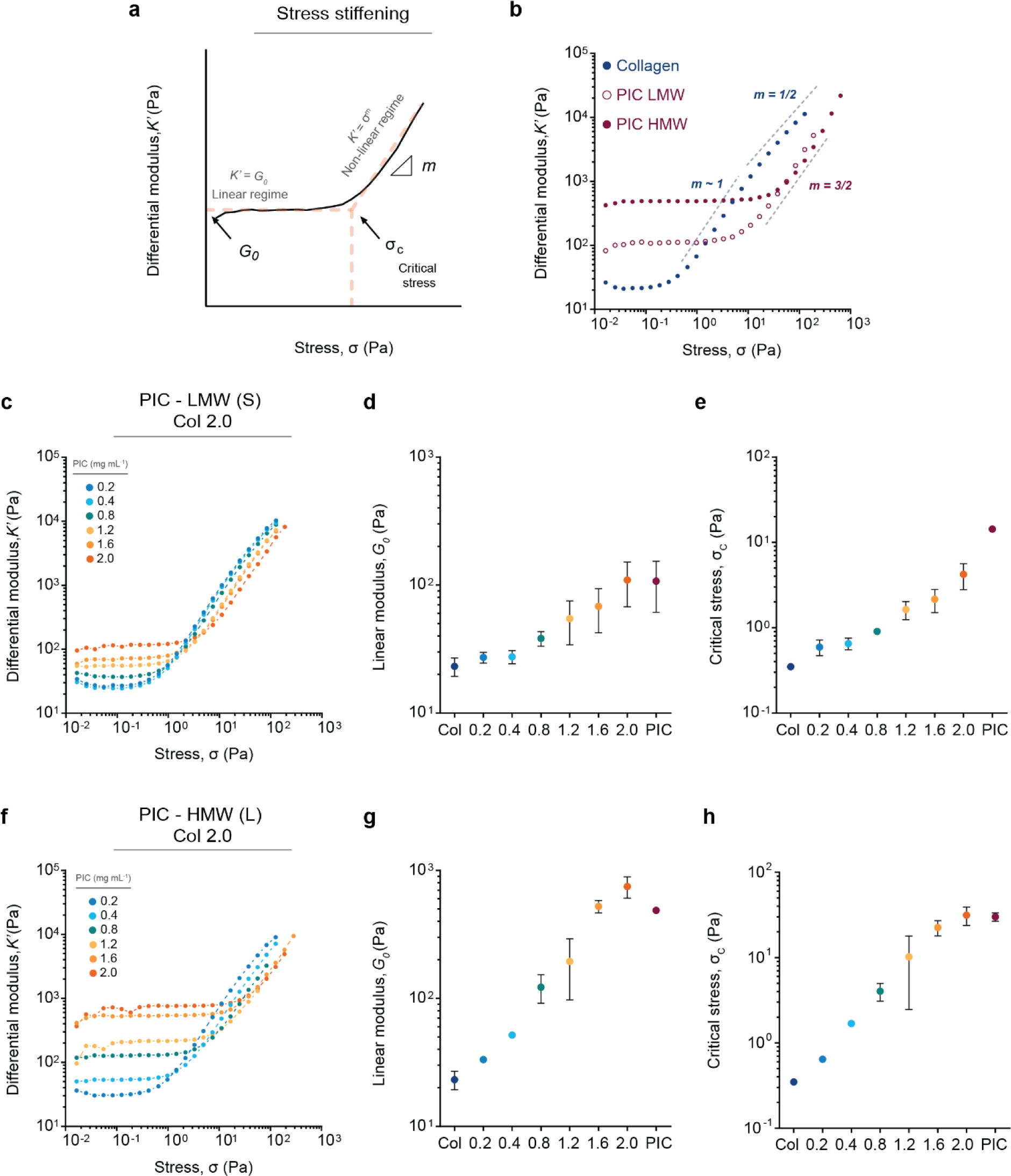
Non-linear mechanical properties of PIC Collagen IPNs. **a**, graphical representation illustrating a non-linear elastic response characteristic of stress-stiffening materials. The response is described by the differential modulus *K’* (*δσ/δγ*) (Pa) as a function of applied stress (*σ*) (Pa). Stress-stiffening materials initially show a linear response at low levels of stress (*K’* = *G_0_*). The material stiffens following *K’* = *σ^m^* (where m is the stiffening index) only when sufficient stress is applied to drive the material into the non-linear regime. This transition is described by the critical stress (*σ_c_*). **b**, non-linear response of pure bovine atelo collagen (2 mg mL^−1^, 1X PBS, pH 7) and PIC hydrogel (2mg mL^−1^, 1X PBS) of LMW and HMW as a function of applied stress. **c**, differential modulus *K’* (Pa) of PIC–Collagen composites containing a constant bovine atelo collagen concentration (2 mg mL^−1^) showing a stress-stiffening response. Increasing the density of the PIC (0.2–2.0 mg mL^−1^) using LMW PIC synergistically increases both the linear elastic modulus (*G_0_*) (**d**) and critical stress (*σ_c_*) (**e**) of bovine atelo collagen. **f-h**, the synergistic mechanical effect is more pronounced when we increase the rigidity of PIC bundles with higher contour length (HMW PIC). Higher amounts of applied of stress are required to deform a collagen network using larger PIC bundles (HMW (L)) compared to shorter PIC bundles (LMW (S)). All data sets show an average of three independent measurements.

### Investigating melanoma behaviour in PIC-Collagen composites

Next, we investigated biological responses of two different cell lines, melanoma, and fibroblasts, to changes in collagen’s non-linear elasticity. As PIC is a biologically inert polymer, this composite system allowed us to investigate the response of cellular behaviour solely to changes in the stress-stiffening response of collagen networks. This offers an advantage over collagen-only hydrogels and other composite systems as it prevents possible effects of changes in collagen concentration, architecture or multivalency experienced when collagen is combined with other biological polymers.

We first investigated morphological responses of a highly motile metastatic melanoma cell line (1205 Lu) (Fig. 4). We embedded melanoma cells for six days in collagen-only and PIC-collagen composites before assessing their morphology (Fig 4a; F-actin and nuclei). Consistent with previous observations,^42^ melanoma cells embedded in collagen-only hydrogels exhibited an elongated morphology with front-rear polarity, extending a singular actin-based pseudopod that allows cells to efficiently translocate within the collagen network.^43,44^ This morphology was conserved in composites with lower PIC concentrations (0.2 mg mL^−1^) while concentrations of ∼ 1mg mL^−1^ and above induced cell clustering (Fig 4a; 0.2 vs. 2 mg mL^−1^). While the number of nuclei increased, cells were still able to form extensions of similar length (Fig. 4 b,c). In addition, we also observe this morphological switch from single elongated cells to multicellular clusters when we embedded cells in collagen composites containing a higher PIC contour length (*L_c_*) (L, HMW) (Fig. 4a; Fig. d,e). Importantly, when comparing the morphological response of melanoma in LMW and HMW PIC-collagen composites, we noted that the clustering of cells was induced at a significantly lower HMW PIC concentration (Fig. 4a; 2.0 mg mL^−1^ LMW PIC, σ_*c*_ 4.21 Pa *cf.* 0.8 mg mL^−1^ HMW PIC, σ_*c*_ 4.03 Pa). While the composites contain different concentrations and contour lengths of PIC, they share a comparable onset of stiffening (σ_*c*_). This indicates that the morphological switching maybe due to cells exerting the same amount of stress against collagen.

**Fig 4.**
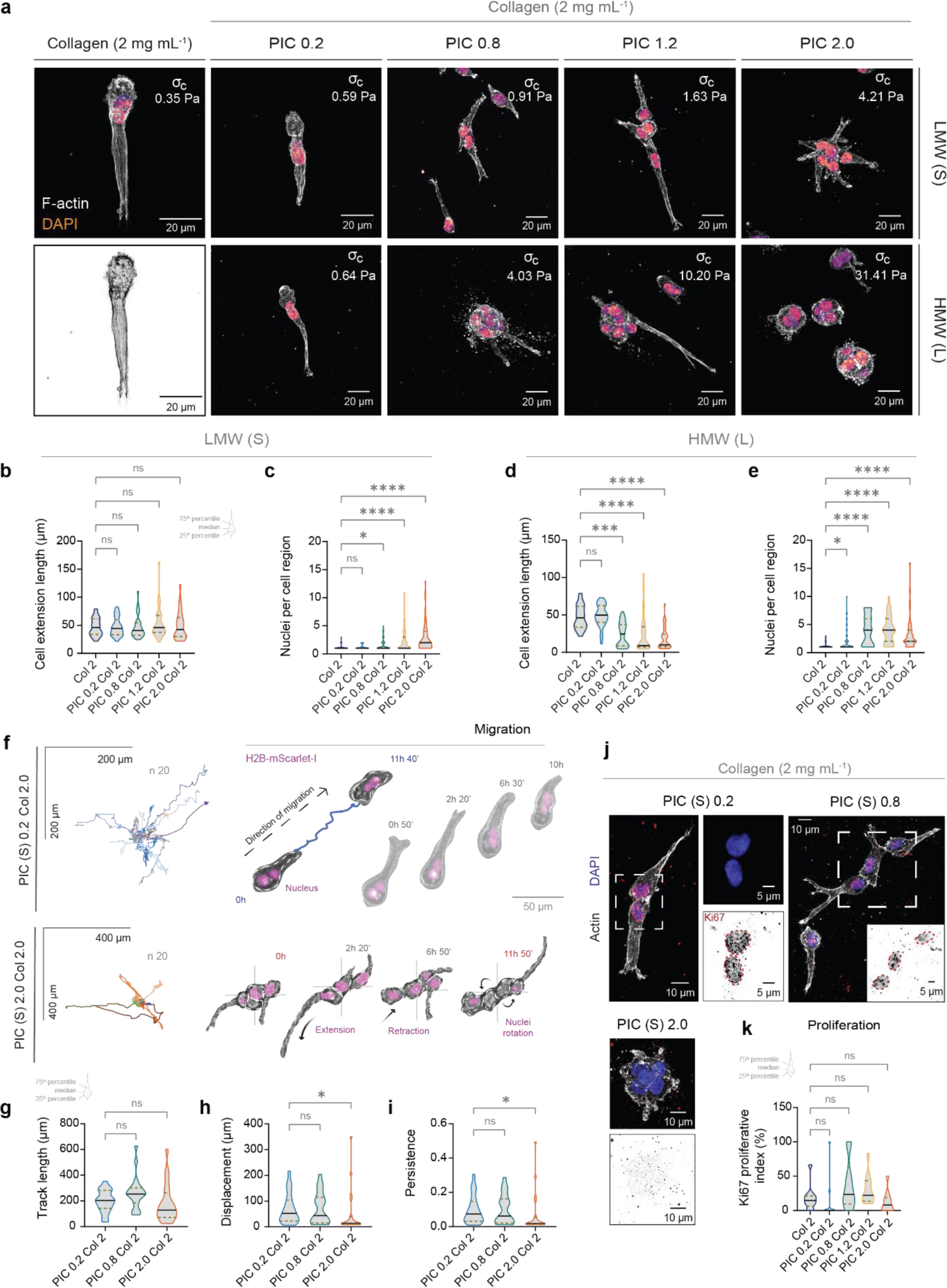
Tuning collagen’s rigidity with PIC modulates melanoma morphology and migration. **a**, confocal images of fixed metastatic melanoma (1205 Lu) cells embedded in pure atelo bovine collagen (2 mg mL^−1^, 1X PBS, pH 7) and LMW and HMW PIC Collagen composites (0.2– 2.0 mg mL^−1^) after six days. **b-e**, quantification of cell extension length (µm) and number of nuclei per cell region. We define the length of a cell extension as the longest distance from the centre of the nucleus to the tip of the pseudopodium. Graph represents the longest extensions in a field of view. Data shows a set of three independent experiments (n = 30 cells per condition). We quantified the number of nuclei within a cell region. A value of one represents an individual cell with a single pseudopodium. Values of 2 and above represent the number of nuclei within one cluster. **d,e** as the concentration of PIC (HMW) increases, there is a decrease in cell extension length and increase in cell nuclei number. Data shows a set of three independent experiments (n = 39–72 cells per condition). Violin plots (truncated, medium smooth) show median (black line) and quartiles (gold pattern lines). **f**, spiderplots of migratory behaviour of melanoma cells in PIC-collagen composites (LMW PIC) with representative time-lapse images. Cells stably expressing the nuclear marker H2B-mScarlet were imaged 72 hours after encapsulation, with images acquired every 10 min for 12 hours. Cells were manually tracked (Image J) using both the centre of the nuclei and perinuclear body in brightfield channels for reference. White dotted line represents cell outline and blue line represents the cell migration path. Time is shown in hours (h) and min (‘). Paths were normalised to the starting position with graphs displaying the 20 longest paths. **g-i**, migratory behaviour is quantified by **g**, the track length (µm), **h**, displacement (µm) and **i**, persistence index ratio in each condition. Violin plots (truncated, medium smooth) show median (black line) and quartiles (gold pattern lines). Data represents three independent biological experiments (n = 20 cell tracks per condition). **j**, representative confocal images of fixed melanoma cells encapsulated in the PIC Collagen composites (LMW PIC) for six days and stained for the proliferation marker, Ki67 (red; magnified Ki67 image shown as contrast inverted image). Ki67 positive cells are shown in composites containing 0.2 mg mL^−1^ and 0.8 mg mL^−1^ PIC. Negative cells are shown in composite containing 2.0 mg mL^−1^ PIC. **k**, the proliferative index (%) for each condition was obtained by quantifying the number of Ki67-positive cells per field of view (five fields of view per sample) over two independent experiments (n = 10 data points per condition). Scale bar (10 µm). Insets (5 µm). Violin plots (truncated, medium smooth) show median (black line) and quartiles (dotted lines).

The morphological switch we observed suggested that the change in collagen mechanics may regulate the ability of cells to migrate. To examine the effect of our PIC-collagen composites on cell migration, we embedded single 1205 Lu melanoma cells and examined their morphology by time-lapse microscopy (Fig. 4 f-i; 12 hours). While cells in all concentrations of LMW PIC were able to form cellular extensions and move within the composites (Fig. 4 f,g), cells in higher PIC concentrations (2 mg mL^−1^) failed to directionally migrate (Fig. h,i). Instead, cells would cluster and rotate, producing short-lived cellular extensions which failed to support the translocation of the cell body (Fig. 4f). In contrast, cells embedded in lower concentrations of PIC were able to efficiently migrate (increased displacement and migration persistence) exhibiting longer-lived pseudopodia (Fig. 4f). This demonstrates that collagen mechanics regulates pseudopodia stability and productive cell migration.

We hypothesised that changes in collagen mechanics may not only induce a migration switch, but may also alter cell proliferation, accounting for the increase in the number of cell nuclei in composites with higher PIC concentration (Fig. 4c; Fig. 4e). To test this, we cultured parental 1205 Lu cells in the PIC-collagen composites for six days, before immunostaining for the cell proliferation marker (Ki67) (Fig. 4j,k). Examination of proliferation (Ki-67-positive nuclei) revealed that cells were able to proliferate in all composites. However, clusters in high concentrations of PIC (2.0 mg mL^−1^) surprisingly lacked Ki67, even though they had the highest number of nuclei (Fig. 4j). Therefore, clusters are likely formed by an early proliferation event after embedding, which ceases once a critical cluster size is met. Together, these results show that increasing collagen’s onset of stiffening (σ_*c*_) controls the ability of cancer cells to migrate and proliferate.

### The onset of collagen’s stress-stiffening response (*σ_c_*) controls fibroblast morphology and collagen remodelling

An important element of pathological conditions, such as wound healing and tumour progression, is the remodelling of collagen by fibroblasts. Thus, we next examined the morphological response of fibroblasts to changes in collagen mechanics using our PIC-collagen composites (Fig. 5; Supplementary Fig. 6; LMW PIC). We correlated collagen remodelling by confocal refection microscopy with changes in fibroblast morphology (F-actin; telomerase immortalised fibroblast, TIF). We found that in lower concentrations of PIC (0.2 mg mL^−1^), cells were able to actively deform the collagen matrix in a similar manner to cells embedded in collagen-only hydrogels (Fig. 5 a,b; Supplementary Fig. 6). However, when we increased the concentration of PIC to 1.2–2.0 mg mL^−1^, cells failed to deform the collagen matrix as indicated by a decrease in collagen reflection intensity at the tips of cellular extensions (Supplementary Fig 6). This was associated with a decrease in cell extension area and circularity (Fig. 5c; Supplementary Fig. 6). Cells became spindle-like in appearance, with long, thin cellular extensions. Together, this indicates that cell-matrix interactions may be adaptively altered in response to increasing the onset of collagen’s stiffening (σ_*c*_) to higher levels of stress.

**Fig 5.**
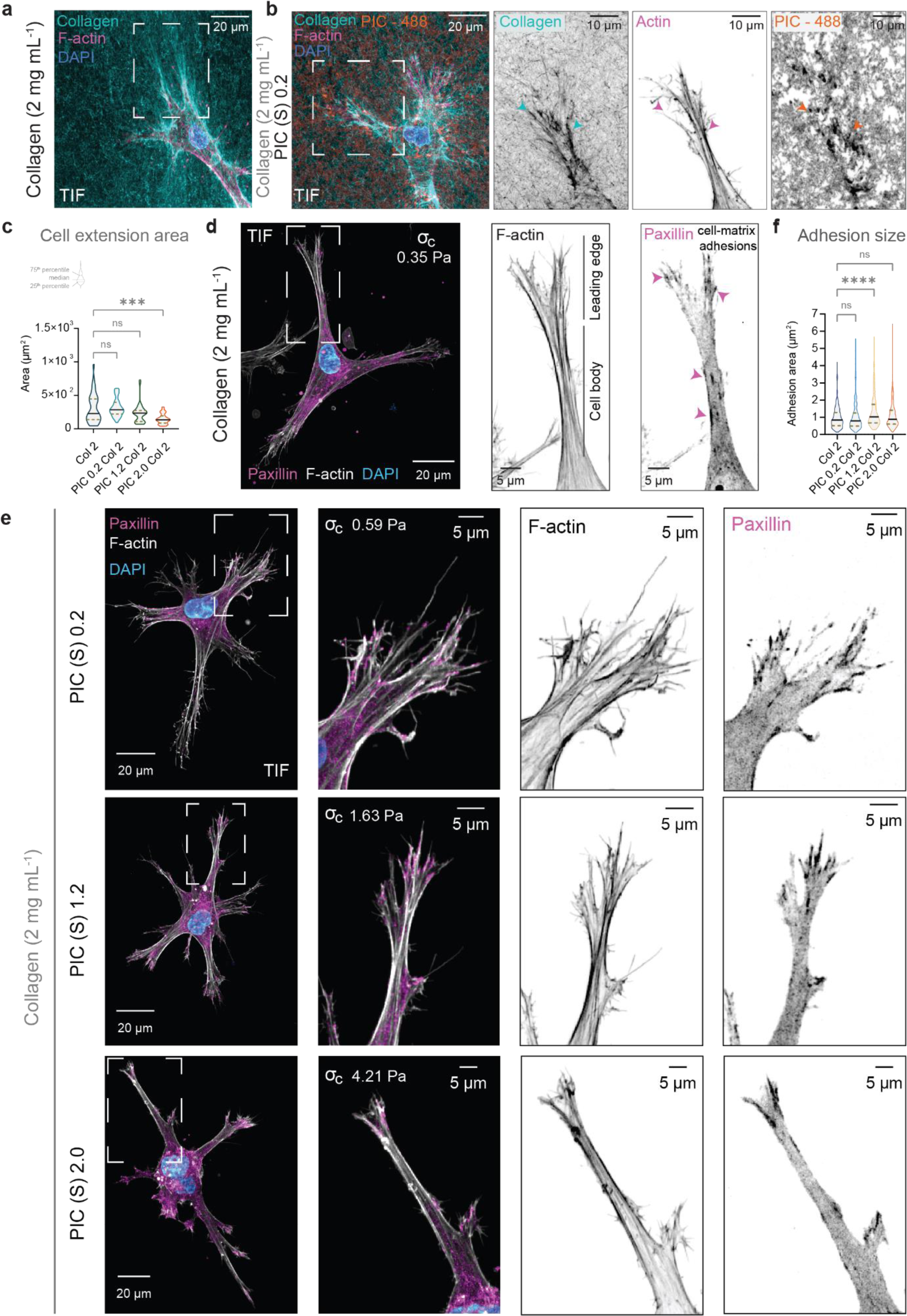
Tuning the onset of collagen’s stiffening (*σ*_*c*_) with PIC stimulates cell-matrix adhesions in 3D collagen hydrogels. **a,b,** super resolution airy scan confocal images of fibroblasts embedded in **a**, atelo bovine collagen (2 mg mL^−1^, 1X PBS, pH 7) and **b**, a PIC–collagen composite (0.2 mg mL^−1^). Samples were fixed 24 hours after embedding. The collagen (cyan) and PIC (orange) networks were directly labelled with a fluorescently labelled collagen-binding protein fragment (I35 mScarlet-i) and PIC (S)-A488, respectively. Actin and nuclei are stained with Phalloidin 647 and DAPI, respectively. Z maximum intensity projections show ∼100 µm. Images were acquired at 37 °C to prevent disassembly of the PIC (S)-A488. Coloured arrows point to regions of increased intensity of both collagen fibres and PIC surrounding the leading edge of the cells indicating simultaneous deformation of collagen and PIC. This indicates that cells are able to interact with collagen fibres in both collagen-only and composite hydrogels. **c**, quantitation of fibroblast cell extension area (µm^2^) in collagen and PIC-collagen composites (0.2-2.0 mg mL^−1^ PIC). Regions of interest (ROIs) of fixed cells were manually acquired in FIJI. Data is representative of two independent biological replicates (n=15 cells per condition). Violin plots (truncated, medium smooth) show median (solid line) and quartiles (dotted lines). **d,** confocal image of a fixed fibroblast expressing Paxillin–mCherry embedded in an atelo bovine collagen hydrogel (2 mg mL^−1^, 1X PBS, pH 7) after 24 hours. Actin and nuclei are stained with Phalloidin 488 and DAPI, respectively. Z projection ∼50 µm. Scale bar (20 µm). Inset shows a zoomed region of a cell extension of interest. The extension is classified into two subregions: leading edge (LE) and cell body (CB). Scale bar (5 µm). **e,** confocal images of fixed fibroblasts expressing Paxillin – mCherry embedded in PIC–Collagen composites (S, LMW; 0.2, 1.2 and 2.0 mg mL^−1^; Collagen 2.0 mg mL^−1^) after 24 hours. Actin and nuclei are stained with Phalloidin 488 and DAPI, respectively. Z projections show ∼50–100 µm depth. Scale bar (20 µm). Insets (5 µm). **f,** adhesion size increases at cell’s leading-edge in response to changes in collagen mechanics. Adhesion size at cell’s leading edge is significantly higher at concentrations of ∼1 mg mL^−1^ PIC. This observation correlates to the concentration at which PIC depicts non-linearity and starts influencing atelo collagen mechanics. Adhesions were manually quantified (n=273–381) from a set of two independent biological replicates (19–25 extensions from 10–14 cells per each condition). Violin plots (truncated, medium smooth) show median (solid line) and quartiles (dotted lines).

### PIC-collagen composites alter cell-matrix interactions in 3D

Cells transmit mechanical forces from their surroundings via focal adhesions. Focal adhesions are multimolecular complexes of proteins that physically connect the rearward flow of intracellular F-actin with the extracellular environment via integrin-transmembrane receptors.^45,46^ It is established that the size and composition of focal adhesion complexes is influenced by the stimulation of tension generated by actomyosin contractility, acting as force-transmitting ‘molecular clutches’ to facilitate forward cell movement.^47,48^ When cells move on glass, focal adhesions grow and elongate when they experience slower actin retrograde flow and tension in the lamella transition zone of the cell leading edge.^49–51^ However, despite this large body of research in two-dimensions, less is known about the spatial organisation of actin and focal adhesions when cells interact with three-dimensional collagen networks.

To investigate the effect of tuning the onset of collagen stiffening (σ_*c*_) on the spatial organisation of actin and focal adhesions, we compared fibroblasts stably expressing the focal adhesion marker Paxillin-mCherry, embedded in collagen-only and composite hydrogels. We chose the adapter focal adhesion protein paxillin because it is present throughout the focal adhesion turnover cycle, at both nascent and maturation stages,^52–54^ ensuring we would capture adhesions. Fibroblasts embedded in collagen-only hydrogels form cell extensions consisting of (1) a peripheral leading edge (LE) which is the tip region, rich in fine actin extensions known as filopodia; and (2) a cell body (CB) which is the pseudopod neck-like region adjacent to the leading edge in front of the nucleus (Fig. 5d; Supplementary Fig. 7). We observed a higher accumulation of actin at lateral regions of the leading edge compared to the central region (Supplementary Fig. 7). Consistent with previous reports,^55^ we also observed that paxillin localises in small puncta at the ends of actin bundles with adhesions that are aligned in the orientation of these actin structures (Supplementary Fig. 7). However, adhesions localised in the neck or lateral region of cell extensions, were elongated (Fig. 5d; Supplementary Fig. 7). This suggests that when cells are embedded in collagen-only hydrogels, cell-matrix interactions at the leading edge are weak, resulting in small punctate adhesions that likely undergo rapid turnover.

In comparison, in the PIC-collagen composites we found that paxillin exhibited a biphasic morphology with a change in spatial distribution in response to decreasing the sensitivity of collagen to stress (Fig. 5e). In collagen composites containing 0.2 mg mL^−1^ PIC, adhesions were small, similarly to those of cells embedded in collagen-only hydrogels (Fig. 5 e,f). However, when we increased the concentration of PIC to 1.2 mg mL^−1^, adhesion size at the tip of the leading-edge increased significantly compared to the cell body. This indicated that cells were exerting tensional forces predominantly at the leading-edge (Fig. 5 e,f; Supplementary Fig. 8). Mechanically, this concentration of PIC corresponds to the point at which collagen starts to lose sensitivity towards stress, suggesting in response, that cells are required to apply more force to deform the collagen matrix resulting in larger adhesions. Increasing the concentration of PIC even further to 2.0 mg mL^−1^ resulted in a decrease in adhesion area at the leading-edge tips (Fig. 5f). This decrease in adhesion size suggests a physical disengagement of the molecular clutch by frictional slippage.^56^ Together, these observations suggest that the onset of collagen stiffening (σ_*c*_) likely modulates the initial force loading rate within focal adhesions in 3D collagen networks. Small increases in σ_*c*_, first results in the stimulation of intracellular tension and growth of strong cell-matrix interactions, but higher tensional forces above a threshold in σ_*c*_, results in weaker interactions with collagen, reducing its deformation.

## Discussion

Our work presents a composite system where the mechanical properties of collagen hydrogels can be easily manipulated with the synthetic polymer, polyisocyanopeptides (PIC). We have shown that, by only introducing small concentrations and different contour lengths (*L_c_*) of PIC, we can influence the mechanical behaviour of collagen hydrogels without changing the reconstitution process of collagen. This system can rapidly undergo gelation by forming a semi flexible background of extremely low synthetic polymer density that mechanically supports the polymerisation of collagen networks, forming an interpenetrating network (IPN). The sequential assembly of this IPN system is a process that its entirely driven by physiological heat (37 °C), making it an optimal system for three-dimensional cell culture.

By combining collagen with polyisocyanopeptides (PIC), we also demonstrate that cells respond to changes in collagen’s non-linear elasticity when collagen experiences mechanical interactions in composite systems. Our findings show that increasing the amount of stress required for collagen to stiffen (onset of stress-stiffening, σ_*c*_) results in cells requiring larger forces to deform collagen networks. This influences biological behaviours in three-dimensions such as cell morphology, migration, growth and the ability of cells to deform their surrounding matrix by cell-matrix interactions.

We first showed that this mechanical phenomenon controls the motility of tumour cells in three-dimensions. Previous reports have shown that cancer cells have the ability to migrate and respond to physical barriers via a switch in migration modes.^57–59^ In particular, melanoma can adaptively move in collagen networks depending on its architecture. In collagen with larger pores, melanoma cells squeeze between fibres using cortical acto-myosin blebbing. However, in a more dense collagen network, melanoma can switch from an ameboid blebbing mode to mesenchymal migration by generating a polarised pseudopodium.^60^ This morphology facilitates the widening of collagen pore sizes by both protease secretion and physical deformations.^44,61,62^ Here, we found that when the collagen architecture is unchanged, but rigidified by synergistic mechanical interactions with a second polymer network, migration becomes non-productive with a morphological switch. Consistent with previous reports, we observed that disruptions in pseudopodia formation decreased persistent migration,^63–65^ resulting in cells forming small clusters. These clusters fail to expand as proliferation ceases when the clusters experience high levels of stress. Importantly, we found that this switch to clustering is dependent on the rigidity of collagen when combined with extremely low concentrations of stiffer PIC polymers. In the context of cell motility, this observation indicates that collagen rigidity independently dictates cell migration in the absence of changes in collagen network density or architecture (i.e. pore size).

In mesenchymal migration, the formation of the pseudopodia in three-dimensional collagen is dependent on focal-adhesion complexes formed by integrin-based interactions.^66^ As these complexes physically link the cellular cytoskeleton to the extracellular environment and mature as a function of force, we hypothesised that increasing the rigidity of collagen networks influences cell–matrix interactions. To this end, we investigated the effect of tuning collagen non-linearity on focal adhesions in fibroblasts. Previous studies have shown that integrin-based adhesions cluster when tuning the rigidity of collagen with different collagen architectures.^67^ However, our results demonstrate that the ability of fibroblasts to actively contract collagen networks is directly influenced by a biphasic response of focal adhesions to the non-linear mechanics of collagen. Consistent with recent insights into how force is transmitted across components of the molecular clutch,^56,68^ we observed that adhesions elongate in collagen hydrogels only at the point at which collagen networks are rigid enough, but still malleable, for cells to actively contract. We entered this mechanical regime, or optimal rigidity, experimentally when combining collagen with ∼1 mg mL^−1^ of LMW PIC. By investigating the spatial organisation of actin and paxillin at the leading edge of cells, we observe that adhesions elongate at the tips where actin bundles accumulate. Increasing the rigidity of collagen with PIC above this mechanical regime results in a decrease in adhesion area, suggesting rapid unbinding events and bond destabilisation within focal adhesions.

We speculate that this biphasic response of focal adhesions may also result in different rates of actin flow with increasing tension at the leading edge, modulating the rate of focal adhesion turnover. Thus, it will be critical for future investigations to determine the spatial localisation of mechanosensitive force transduction proteins such as vinculin and talin. Key to this will be gentle, advanced volumetric imaging techniques capable of achieving subcellular resolution of focal adhesion turnover dynamics in 3D.^69^ Ultimately, this will couple our understanding of the dynamics of the molecular clutch engagement at the different stages of stress stiffening, in materials exhibiting non-linear elasticity.

Our results also show that cells exhibit much richer responses when collagen experiences mechanical interactions with other polymers. Therefore, further field application of our composite system will facilitate the exploration of morphological responses that are not commonly observed in collagen only hydrogels, as well as the molecular mechanisms that underpin them. In the context of tissue homeostasis, our findings provide deeper insight into how cells respond to an increase in tension as they encounter larger depositions of matrix during wound healing and pathological conditions such as cancer.

## Methods

### Synthesis of Polyisocyanopeptides (LMW, HMW, PIC-N_3_, PIC-Alexa488)

Following a previously described procedure for the synthesis of polyisocyanopeptides,^31–33^ polymers containing methoxy-functionalised tri-ethylene glycol (EG_3_) monomers were synthesised using 1:1,000 (Low molecular weight, LMW, S – short polymer) and 1:10,000 (High molecular weight, HMW, L – long polymer) catalyst/monomer ratios. Polymers were characterised by viscometry (LMW M_v_: 283 kg mol^−1^, HMW M_v_: 518 kg mol^−1^) and rheology in 1X Phosphate Buffer Solution (PBS). LMW PIC-Azide polymers were synthesised as described using an EG_3_-Azide functional monomer in a 1:100 Azide: Methoxy ratio (Mv: 170 kg/mol). LMW PIC-Azide was reacted with AlexaDye 488 – DBCO as previously reported with minor modifications.^39^ LMW PIC-Azide (100 mg) was dissolved in 50 mL of ACN for 48 hours at room temperature. Then, a solution of AlexaDye 488 DBCO (3.123 x10^−7^ mol, 0.3108 mg dye) from a stock solution (2.99 mM, DMF) was added to the PIC solution and stirred for 48 hours at room temperature while avoiding light exposure. The polymer was precipitated in cold diisopropyl ether, purified by dialysis (MWCO 14 kDa) at 4 °C for 24 hours against MQ water and recovered by lyophilisation.

### Preparation of collagen solutions

All reagents were cooled on ice before use. Collagen Type I atelo of bovine origin (Nutragen^®^, 5.9 mg mL^−1^ in 0.01 M HCI, #5010, Advanced Biomatrix) was diluted with 1-part 10X PBS, followed by neutralisation to pH 7 with 0.1 M NaOH and diluted with distilled MQ water to generate solutions of 4 mg mL^−1^. Supplementary methods Table 3.3.

### General preparation of PIC–Collagen composites

All experiments were performed using the methoxy-functionalised tri-ethylene glycol (EG_3_) PIC polymers unless stated otherwise. To form the PIC–Collagen composites, PIC of LMW or HMW was dissolved in 1X PBS to generate a stock solution of 4 mg mL^−1^. The PIC solutions were dissolved in an automatic rotator at 20 rpm for 72 hours at 4 ^°^C. The stock solution of PIC was diluted in a series of dilutions with cold 1X PBS to produce solutions of 200 µL of final volume. Supplementary methods Table 3.4. Next, 200 µL of the prepared PIC solution (0.4, 0.8, 2.4, 3.2, 4.0 mg mL^−1^) was added to 200 µL of a cold solution of collagen (4 mg mL^−1^, 1X PBS) to give composites with final PIC concentrations of 0.2, 0.4, 0.8, 1.2, 1.6 and 2.0 mg mL^−1^ while containing a final collagen concentration of 2.0 mg mL^−1^. The composites were mixed by pipetting while swirling to avoid bubbles and kept on ice prior to their characterisation.

### Rheology

Collagen gels (2 mg mL^−1^) and PIC (0.2–2.0 mg mL^−1^) – Collagen (2 mg mL^−1^) composites were analysed using an Anton Paar MCR-502 WESP rheometer. Samples were transferred to a pre-cooled (5 °C) bottom stainless-steel plate (Anton Paar) setting a gap of 0.5 mm using a stainless-steel parallel plate PP-25 (25 mm diameter, Anton Paar). Samples were trimmed and a thin layer of silica oil was placed around the sample. Gelation was induced with a temperature ramp from 5 °C to 37 °C (1 °C min^−1^) and further incubated for 2 hours at 37 °C. Changes in viscoelastic properties were monitored by applying oscillatory shear rheology at a constant strain of 0.5 % and frequency of 1 Hz. A frequency sweep from 10 to 0.1 Hz at constant strain of 0.5% was then applied and the non-linear profile was evaluated using previous pre-stress protocol.^31^ The linear elastic modulus *G_0_* represents an average of the first three individual values in the linear regime at low levels of stress. Following previous studies, ^32,33^ we obtained the onset of stiffening (*σ_0_*) by performing a linear interpolation of the differential modulus *K’* (*δσ/δγ*) curve for each condition and determined the minimal stress values.

### Confocal Rheology

Simultaneous confocal imaging and rheology was performed with a custom built Confocal– Rheometer by manually sliding a rheometer with an open bottom plate configuration (MCR-502 WESP rheometer, Anton Paar) placed on a rail stage until positing on top of an inverted confocal microscope (TCS SP8 WLL, Leica Microsystems).^70,71^ All measurements were performed with oscillatory shear rheology and a measuring stainless steel parallel plate PP-25 (25 mm, Anton Paar). An extension tube was used for installing the objectives under the rheometer stage. For assembling the rheometer sample holder, in general, a 3 mm plastic ring was placed under a thin steel plate with objective hole. A glass cover slip (50 mm, # 1.5) was placed on top of the thin steel plate and sealed with sealant ring (rubber) and stainless streel ring with rubber ring.

#### Confocal radial distance calibration

The radial distance of the objective was calibrated within ∼ 1 mm form the centre of the plate in order to reduce visualised motion from the oscillation. To calibrate, scans of 512 x 512 pixels (465 x 465 µm) were acquired with a 20x Air objective using reflection mode and gap between plates of 0.1 mm. A 488 nm laser with an emission range 479–498 nm using a PMT detector were used to acquire the reflection from the bottom surface of the plate. A steady shear rate of 0.3330 (s^−1^) was applied while simultaneously imaging at a rate of 27.87 (s^−1^) for 30 seconds. Distinguishable defects on the surface of the plate were used to calculate the radial position of the objective by calculating the local linear velocity.

#### Image acquisition

After calibration, a HC PL APO CS2 40x/1.10 water-immersion objective (Leica Microsystems) with correction collar of 0.18 was placed under the lower plate and glass cover slips were pre-cooled to 5 °C with a gap of 0.5 mm for 10 min before placing each sample. After placing the samples, the measuring gap was set the 0.5 mm and sample were trimmed and a thin layer of silica oil was placed around to avoid dehydration. Fields of views were set to at least ∼50 µm above the bottom glass. The reflection intensity of the collagen fibres was acquired using a 550 nm laser and emission range 530–570 nm with a PMT detector. Stacks of 20 µm size were acquired with a step size of 1.0 µm every 1 min for 2.5 hours. All scans were 1024 x 1024 pixels (61.44 µm x 61.44 µm) and acquired in a bidirectional scan direction with a line accumulation of 2. Time zero is defined as the time when the starting temperature is 5 °C. The general protocol for inducing gelation with temperature ramp (5– 37°C, 1 °C min^−1^) and incubation for 2 hours at 37 °C was applied and changes of viscoelastic properties (*G’* and *G’’*) were monitored over time.

#### Co-imaging of fluorescence and reflection modes

To acquire both channels at the same rate as the temperature increase (1 °C min^−1^), stacks of 10 µm size were acquired with a step size of 1.0 µm every 1 min for 2.5 hours. The fluorescence intensity of the PIC-A488 was acquired using a 488 nm laser and emission range 510-560 nm. The reflection intensity of the collagen fibres was acquired using a 650 nm laser (%) and emission range 630–570 nm with a PMT detector. Scans were acquired sequentially between frames while collecting the reflection of the fibres in the first line. Stacks of 20 µm size were acquired with a step size of 1.0 µm every 1 min for 2.5 hours. All scans were 1024 x 1024 pixels (61.44 µm x 61.44 µm) and acquired in a bidirectional scan direction with a line accumulation of 2. Time zero is defined as the time when the starting temperature is 5 °C. After gelation for 2 hours, images of 50 µm depth from three different regions were acquired at 37 °C using this set up to maintain the formation of the PIC network. Maximal projections were processed in FIJI and three-dimensional reconstructions were processed in Imaris x 64 (version 9.7.2).

### Pore analysis by confocal laser scan microscopy

Collagen and PIC–Collagen composites (1X PBS) were imaged by confocal reflection microscopy (CRM) using an inverted confocal microscope (TCS SP8 WLL, Leica Microsystems). Hydrogels were incubated at 37 °C for 2 hours in a 15-well chamber slide (ibidi, USA) and heating block. Samples were imaged at room temperature (∼21 °C) after gelation. A 488 nm laser and emission range of 478–498 nm using a HC PL APO CS2 40x/1.10 water-immersion objective (Leica Microsystems) with correction collar of 0.18 and HYD detector were used to acquire data. To evaluate homogeneity throughout the samples, scans were acquired ∼50 µm from the bottom layer and three consecutive stacks of 50 µm size were acquired with a step size of 1.0 µm. All scans were 1024X1024 pixels (235×235 µm). Maximal projections were generated in FIJI by applying a mean and gaussian blur filter filtering out features smaller than one-pixel before binarization. An automated analysis was used to determine the pore size of the collagen networks using open-source MATLAB algorithm. ^72,73^

### Small Angle Neutron Scattering – Ultra Small Angle Neutron Scattering (SANS-USANS)

Combined small-angle neutron scattering and ultra-small angle neutron scattering (SANS/USANS) techniques were utilized to determine the structural properties of the collagen networks with a linear increase in PIC concentrations. Both small-angle neutron scattering (SANS, QUOKKA)^74^ and ultra-small angle neutron scattering (USANS, KOOKABURRA)^75^ were performed at Australian Nuclear Science and Technology Organisation (ANSTO, Sydney, Australia).

#### Sample preparation

All neutron scattering experiments were performed using the methoxy-functionalised tri-ethylene glycol (EG_3_) PIC polymers of LMW (M_v_: 283 kg mol^−1^) in 1X PBS. Collagen solutions using bovine Atelo Type I Collagen (Advanced Biomatrix, Nutragen^®^, 5.9 mg mL^−1^) were prepared at 4 mg mL^−1^ in 1X PBS. For SANS (QUOKKA) experiments, we prepared 500 µL of sample while for Kookaburra we prepared 2.0 mL of sample.

#### SANS-USANS Acquisition

All SANS and USANS measurements were performed at 37 °C. For SANS, a whole q range of 7×10^−4^ to 0.1 Å^−1^ was covered using an incident wavelength of 6 Å with 10% resolution. Three instrument configurations were conducted including source-to-sample (SSD) = sample-to-detector distances (SDD) = 20 m, SSD = SDD = 8 m, and SSD = 4 m while SDD = 1.3 m. The source and sample aperture diameters were 50 mm and 12.5 mm, respectively. Samples were loaded in quartz cuvettes with 2 mm thickness were measured. For USANS, samples were measured in a double-crystal diffractometer using a Gd aperture with a diameter of 29 mm and a neutron wavelength of 4.74 Å. The q range of 4×10^−5^ to 8×10^−4^ Å^−1^ was covered. SANS data were then reduced and merged with desmeared USANS data as previously described^76–78^. Combined SANS/USANS scattering data were then plotted. Scattering curve fitting and analysis were conducted on SASView 5.0.5. For the structural analysis of fiber diameter, the scattering data probed in the high q range (SANS) were analyzed using a Guinier-Porod model following Equations 1, where I(Q) is the scattering intensity at scattering vector (Q) and Rg is the radius of gyration.

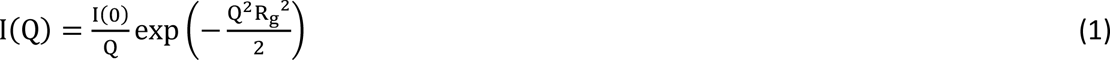

Similarly, the pore sizes of the networks were analyzed by the same model using the scattering data in the USANS range. Average fiber diameter (D) or average pore size (ξ) was calculated using Equation 2.

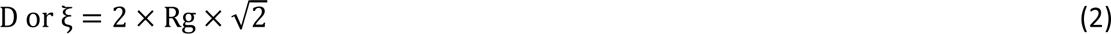

#### Contrast variation

To separate the neutron scattering intensity of PIC from the collagen in the PIC-Collagen composites, we first performed a contrast variation experiment using SANS (QUOKKA) and determine the match point of the PIC. To this end, we measured the neutron scattering of a series of solutions where we linearly increase the concentration of deuterium oxide (D_2_O) in PIC solutions of 2 mg mL^−1^. We prepared the solutions with different ratios of D_2_O/H_2_O (0, 20%, 40%, 50%, 60%, 80% and 100% D_2_O) using ratios of two 100.0 mL solutions of 1X PBS (one solution using 100% MQ water and another using 100% D_2_O). We observed that the match point of PIC was at 20% D_2_O. Once we determined the match point, we prepared stocks of 2 and 4 mg mL^−1^ of PIC by dissolving the polymers in a solution of 40% D_2_O containing 1X PBS in an automatic rotator at 20 rpm for 72 hours at 4 ^°^C. On site, we diluted the stock solutions using a solution containing 40% D_2_O/1X PBS to generate solutions of 0.4, 2.4 and 4.0 mg mL^−1^ PIC (1X PBS) and combined equal amounts of volume to a solution of collagen prepared with 100% MQ using 1X PBS as previously prepared for the other experiments. Note that for the contrast variation experiments requiring higher percentages of D_2_O, we prepared a stock solution of PIC in 100% D_2_O/1X PBS and dialysed the collagen stock solution in 100% D_2_O. For the dialysis, we transferred ∼5 mL collagen solution to a dialysis tubing membrane (SnakeSkin™, MWCO 7,000, 22 mm x 35 ft dry diameter) and dialysed against 50 mL of 100% D_2_O with 0.02 M acetic acid in a falcon tube overnight at 4 °C. We observed no precipitation when recovering the collagen from the membrane. To reconstitute the collagen in 90 % D_2_O, we added the1 part of 10X PBS (D_2_O) after mixing the collagen stock first with the PIC solution (100% D2O, 1X PBS), 0.1 M NaOH and the corresponding amount of D_2_O to make up the final volume.

### General cell culture

Human melanoma cell lines (1205 Lu) and Telomerase-immortalised fibroblasts (TIF) were cultured in T75 cell culture flasks with 10.0 mL of a general cell culture media solution of 500 mL Dulbecco’s Modified Eagle Medium (DMEM, high glucose, pyruvate, no glutamine; Gibco, 10313021) containing 50 mL of 10% of foetal bovine serum (FBS; Gibco, 10100147), 5 mL Penicillin-Streptomycin (10,000 U/mL; Gibco), and 5 mL Minimum Essential Medium Non-Essential Amino Acids (MEM NEAA, 100 X, Gibco). 1205 Lu endogenously labelled for meGFP-α-Tubulin (CRISP) expressing mScarlet-I H2B^42^ and TIFs expressing Paxillin-mCherry was established by lentiviral transduction.^54,79^ TIF cells were cultured in 10 µg/mL Blasticidin S Hydrochloride (Gibco, A1113903).

### General procedure for cell embedding in PIC-Collagen composites

All collagen and PIC solutions were prepared following the sample preparation used for rheology experiments using *sterile* solutions of PBS (10X, 1X) and MQ water. Note that for cell culture experiments we used 12 µL a sterile solution of 7.5% sodium bicarbonate (Gibco) to neutralise the collagen solution to pH 7. We adjusted the volume with 32.4 µL of sterile water to make a final volume of 200 µL. The amount of 10X PBS was kept consistent (Supplementary methods, Table 4.1). All reagents were kept in ice before embedding. In general, cells were detached and resuspended in 1 mL of media. A cell pellet of 5.0 x10^4^ cell mL^−1^ was generated by quickly centrifuging the solution containing the cells in 1.5 mL Eppendorf tube at high speed for 10 seconds. Media was removed slowly without disturbing the cell pellet. The cold collagen solution (4 mg mL^−1^, 1X PBS, pH 7) was added on top of the pellet and mixed by pipetting while swirling. The solution was placed momentarily in ice to prevent polymerisation. 40 µL of the collagen solution containing the cells were mixed by pipetting with 40 µL of the desired PIC solution in a separate 1.5 mL Eppendorf tube. 30 µL of the material/cell mixture were transferred to a well from a pre-heated (37 °C) µ-Slide 18 Well Glass Bottom dish (# 81817, Ibidi). The samples were incubated for at least 2 hours in a humidified chamber at 37 °C/ 5% CO_2_. 70 µL of general cell culture DMEM containing 20 mM HEPES were added on top of hydrogels and incubated at 37 C/5% CO_2_ before imaging or fixation.

### General procedure for fixation

All cells were fixed by removing the cell media and adding 70 µL a solution of 4% Paraformaldehyde (PFA) in 1X Brinkley Buffer 1980 (5X BRB80; 400 mM PIPES, 5 mM MgCl_2_, 5 mM EGTA, pH 6.8) and MQ water for 15 min at room temperature. The fixation solution was removed, and the samples were washed twice with 70 µL of 1X PBS+ (0.9 mM Ca^2+^ and 0.5 mM Mg^2+^) for 5 min. 70 µL of 1X PBS were added to the samples before staining or for storage at 4 °C.

### 1205 Lu in PIC-Collagen composites

#### Live cell imaging experiments

Cells were imaged by live cell imaging two hours post encapsulation. Imaging was performed in an Andor Dragonfly Spinning Disc Scanhead confocal microscope with dual Andor Zyla 4.2 sCMOS cameras equipped with CO_2_ and temperature-controlled chamber (37 °C) and a LED Cool/LED pe-100 illumination system. Time-lapse imaging was acquired with a 20x Dry objective (Plan Apo VC, 0.75 N.A.) by sequentially acquiring fluorescent channels 488 mm (30%, 525–550 nm: 200 ms, confocal 25 µm), 561 mm, (20%, 600-50 nm: 200 ms, confocal pin hole 25 µm) and brightfield channels (15% intensity LED Cool/LED pE-100, 400 ms, confocal pin hole 25 µm). Scans were acquired every 10 min for 12 hours using a fully monotonised X-Y-Z stage capturing scans in at least two different regions per sample. Cell tracking analysis was performed using the Manual Tracking plugin in FIJI to determine the track length (µm), displacement (µm) and persistence (i.u) travelled in each condition.

#### Proliferation experiments

1205 Lu parentals were embedded in collagen and PIC-Collagen composites for 6 days and fixed. After fixation, cells were permeabilised with 70 µL of 0.5 % Tx-100/PBS for 10 min at 4 °C and blocked in 70 µL of a solution containing PBS+, saponin and fish skin gelatin (PFS) (500 mL PBS+, 3.5 g fish skin gelatin, 1.25 mL 10% saponin/PBS, 0.02% NaN_3_) for 1 hour at room temperature while gently rocking. The primary antibody (rabbit polyclonal to Ki67 (1:250): ab15580, abcam) was diluted in PFS. 50 µL of the antibody solution was added to each well and incubated for 2 hours at room temperature while gentle rocking. Samples were washed three times by incubating 70 µL of PFS for 5 min at room temperature while gently rocking. Next, the secondary antibody Janelia Fluor 549 (polyclonal anti-rabbit IgG (H+L), 1:200, NBP1-75286JF549, Novus Biologicals) and Alexa Fluor™ 488 Phalloidin (1:500, A12379, Invitrogen™) were diluted in PFS. 70 µL of the solution was added to each well and incubated for 2 hours at room temperature while gently rocking and avoiding light exposure. Samples were washed three times by incubating 70 µL of PFS for 5 min at room temperature while gently rocking. 70 µL of a DAPI solution in 1X PBS (1:10,000, D1306, Invitrogen™) were added to each well and incubated for 10 min at room temperature while gently rocking and avoiding light exposure. Samples were washed with 70 µL 1X PBS for 5 min at room temperature while gently rocking and stored with 70 µL 1X PBS at 4 °C. Imaging was performed in an Andor Dragonfly Spinning Disc Scanhead microscope with a 40x water objective (Apo λ LS, 1.15 N.A.) using GenTeal™ gel (Alcon Laboratories) for immersion to avoid rapid evaporation of water. Images were acquired sequentially in the following order: 561 nm (50%, 600-50 nm: 400 ms. confocal 25 µm), 488 nm (10%, 525–550 nm: 200 ms, confocal 25 µm), 405 nm (20%, 450-50 nm: 200 ms, confocal 25 µm). Stacks of 50-70 µm size were acquired with a step size of 0.2 µm while acquiring all Z positions for each channel. At least five different regions were acquired for each condition. All scans were 2048 x 2048 pixels (16-bit), binning 1X1 with a frame averaging of 1.

#### Image processing

After acquisition, images were maximally projected in FIJI and deconvolved using Microvolution™ Deconvolution (Version 2016.02). In general, 10 deconvolution iterations were run per channel and background was corrected at 1%. A confocal PSF model was specified. The vectorial model was used as the mathematical approximation of microscope diffraction. A pinhole spacing of 625 nm and back projected pinhole radius of 312.5 nm was specified for all emission wavelengths.

### Human dermal fibroblasts in PIC-Collagen composites

#### Cell-matrix adhesion experiments

TIF cells expressing Paxillin-mCherry were embedded in collagen and PIC-Collagen composites for 24 hours before fixation. Alexa Fluor™ 488 Phalloidin (1:500, A12379, Invitrogen™) and DAPI (1:10,000, D1306, Invitrogen™) were diluted in 1X PBS directly added to the sample without permeabilization. Samples were incubated for 2 hours at room temperature while gently rocking before washing three times with 1X PBS as previously reported. Imaging was performed in an Andor Dragonfly Spinning Disc Scanhead microscope with a 40x water objective (Apo λ LS, 1.15 N.A.) using GenTeal™ gel (Alcon Laboratories). Images were acquired sequentially in the following order: 561 nm (70%, 600-50 nm: 400 ms. confocal 25 µm), 488 nm (10%, 525-50 nm: 200 ms, confocal 25 µm), 405 nm (20%, 450-50 nm: 200 ms, confocal 25 µm). Stacks of 100-150 µm size were acquired with a step size of 0.2 µm while acquiring all Z positions for each channel. At least five different regions were acquired for each condition. All scans were 2048 x 2048 pixels, 16-bit, binning 1X1 with a frame averaging of 1. Images were deconvolved as reported above.

#### Morphology and Paxillin-mCherry analysis

Deconvolved images were processed in FIJI by adjusting the minimum and maximum sliders of each channel for presentation purposes. No other image processing was performed.

#### Collagen remodelling experiments

For the images showing the remodelling of collagen with PIC-A488, cells were embedded in PIC(S) Alexa-488 at 0.2 mg mL^−1^ (1X PBS). After fixation, cells were stained with Alexa Fluor™ 647 Phalloidin (1:250, A22287, Invitrogen™), DAPI (1:10,000, D1306, Invitrogen™) and recombinant mScarlet-I CNA35 collagen I binding peptide^42^(1:500) in 1X PBS. Images were acquired in an inverted LSM880 Fast Airyscan (Zeiss) using a Plan Apochromat 40X water objective (1.2 N.A.) using GenTeal™ gel. Acquisition was performed sequentially in the following order 633 nm (605-70 nm), 561 nm (605-70 nm), 488 nm (525-50 nm), 405 nm (445-50 nm) using a 3.0% laser power with a gain 800% and digital gain 1.0 for all channels. Scans were 1052×1052 µm (16-bit) and acquired in a bidirectional scan direction with a speed of 6 and line averaging of 1. Stacks of 100-150 µm were acquired using a 0.22 µm step size while acquiring all Z positions for each channel. Airy scan images were processed in the ZEISS software (Zeiss Zen 2012 Black). For the images showing the remodelling of collagen in reflection mode, TIFF parentals were embedded in collagen and PIC-Collagen composites for 24 hours before fixation. Cells were stained with Alexa Fluor™ 488 Phalloidin (1:500) and DAPI (1:10,000, D1306, Invitrogen™) in 1X PBS as reported. Images were acquired in a inverted confocal microscope (TCS SP8 WILL, Leica Microsystems) using HC PL APO CS2 40x/1.10 water-immersion objective (Leica Microsystems).

### Software and statistical analysis

Statistical significance was determined in GraphPad Prism (9.4.1) for all sets of data by first identifying outliner using the ROUT (Q=1%) method and applying a Normal (Gaussian) distribution test using a Shapiro-Wilk test distribution. We applied a nonparametric test using a Kruskal-Wallis one-way test and Dunn’s multiple comparison tests to assess significance. Data reports confidence level of 0.05. Analysis and graphs include outliers identified. Adobe Illustrator 2022 (Adobe) was used to generate all figures.

## Author contributions

M.A.E.M conceptualised and directed the project. M.A.E.M designed, performed, and analysed all rheological, microscopy and biological experiments with guidance of J.L and S.J.S. M.A.E.M prepared figures and wrote the original draft. Z.W performed and analysed neutron scattering experiments. R.J.J contributed cell lines and provided biological reagents. P.T designed the confocal-rheology set up, provided script and guidance with pore size analysis and confocal rheology. J.M and E.P.G designed and performed neutron scattering experiments. J.L provided chemical reagents and provided supervision in rheological experiments. S.J.S provided access to instrumentation, biological reagents and provided supervision in biological experiments. A.E.R provided access to instrumentation and conceived the project. S.J.S and A.E.R Funding acquisition. M.A.E.M and S.J.S edited the manuscript.

## Data and materials availability

All data are available in the main text or supplementary materials.

## Supplementary

**Supplementary Fig 1.**
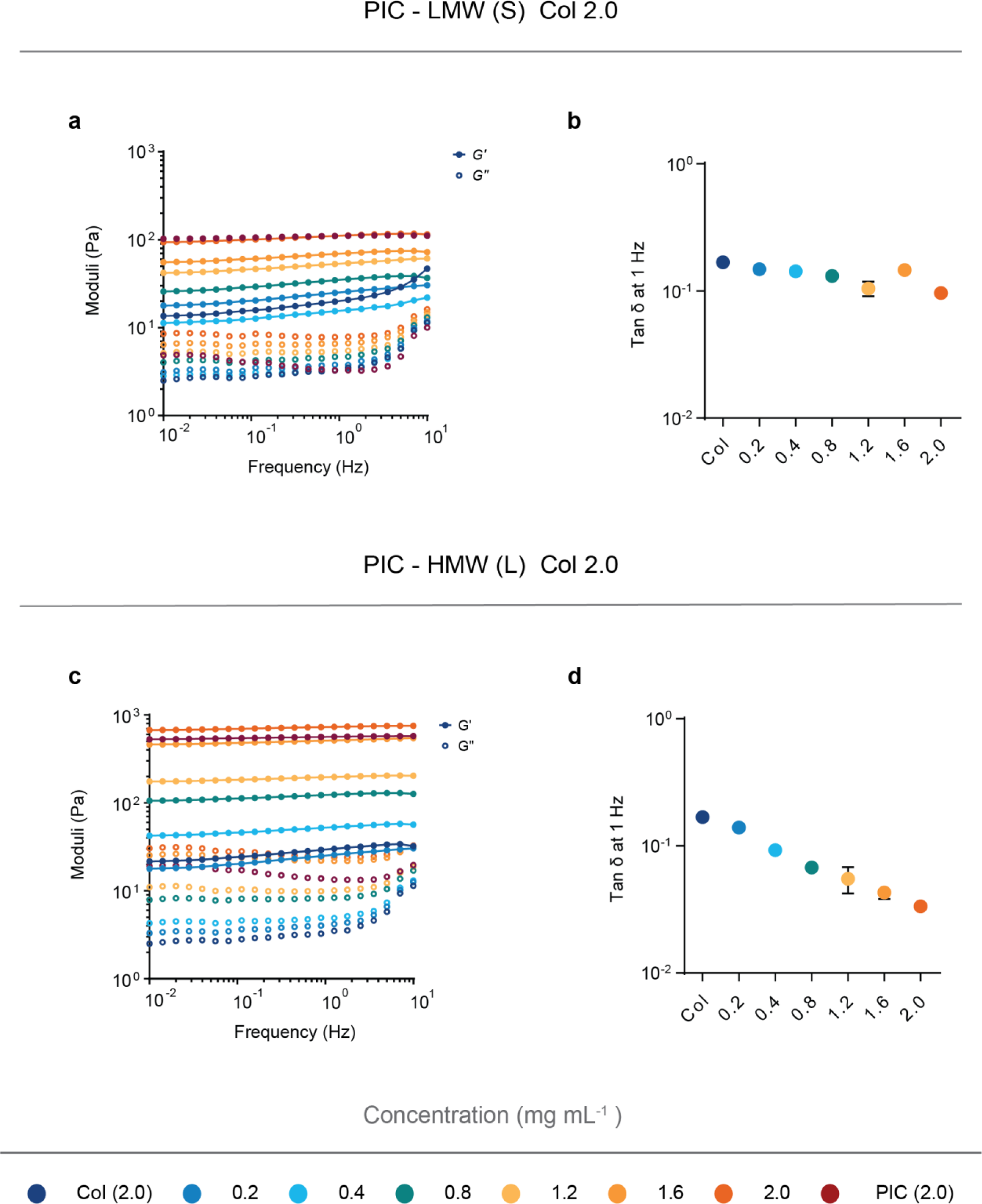
Extended linear mechanical characterisation of PIC Collagen composites. **a-d**, the viscoelastic properties of the composite hydrogels were evaluated after gelation with applied frequency sweeps (10 - 0.01 Hz). **b,d**, The *tan δ* (*G’’*/*G’*) profiles indicate that the viscoelastic profile of bovine atelo collagen is consistent throughout all the PIC-Collagen composites containing lower molecular weight (**a,b**), but the elasticity of the materials increases with an increase in PIC contour length (**c,d**). All sets of data represent three-independent measurements.

**Supplementary Fig 2.**
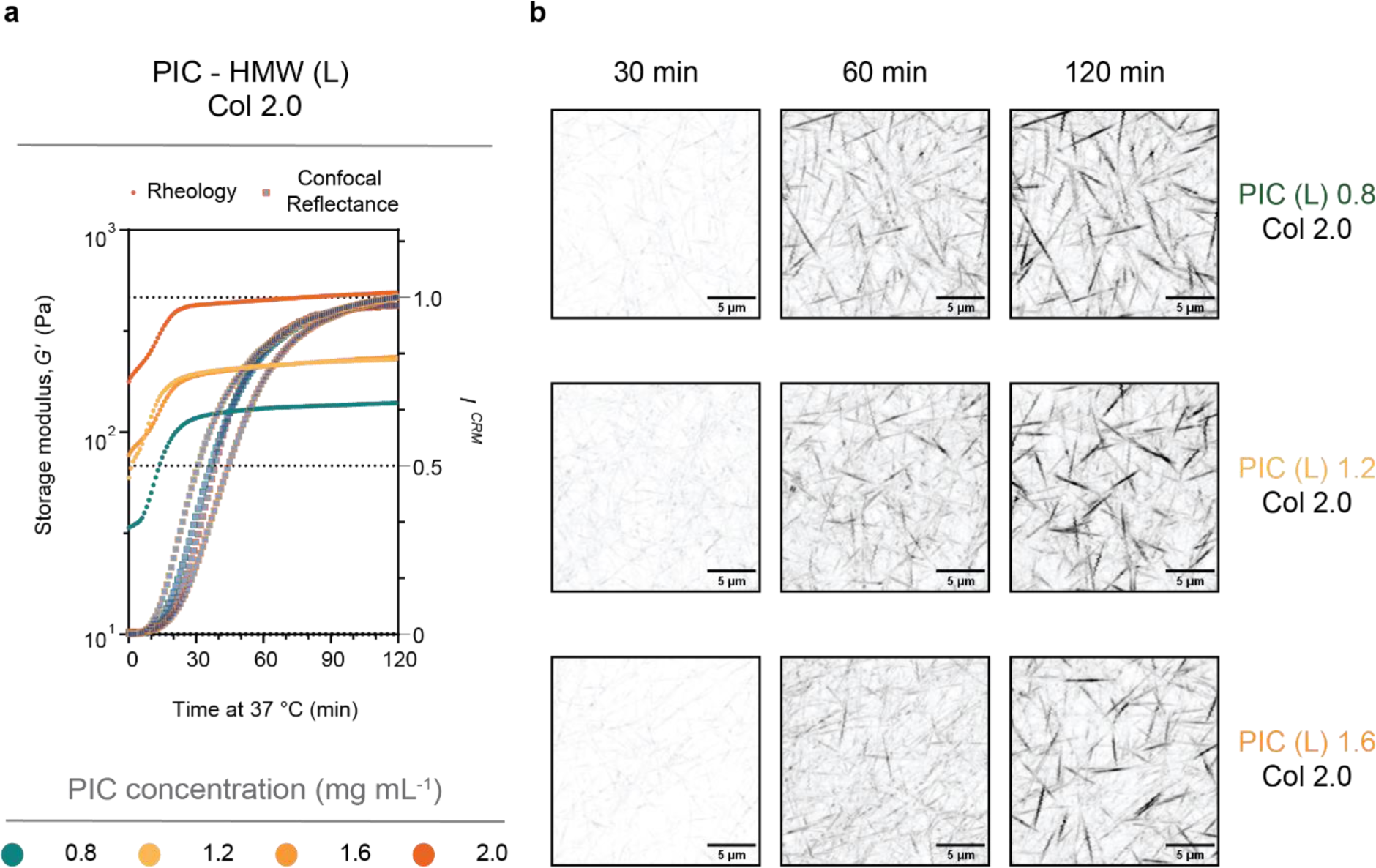
Monitoring collagen fibrillogenesis with increasing PIC density using confocal rheology. **a-b**, the sol–gel transition of the PIC-Collagen hybrids occurs before the increase in *I^CRM^*. Interestingly, all composites show an identical *I^CRM^* profile indicating that the intensity profile collagen network formation is not accelerated with increasing PIC polymer density (**a**). This can also be observed by inverted confocal images (**b**). Z-projections show 20 µm depth. Scale bar (5 µm). Data represents an average of two independently prepared samples for each condition.

**Supplementary Fig 3.**
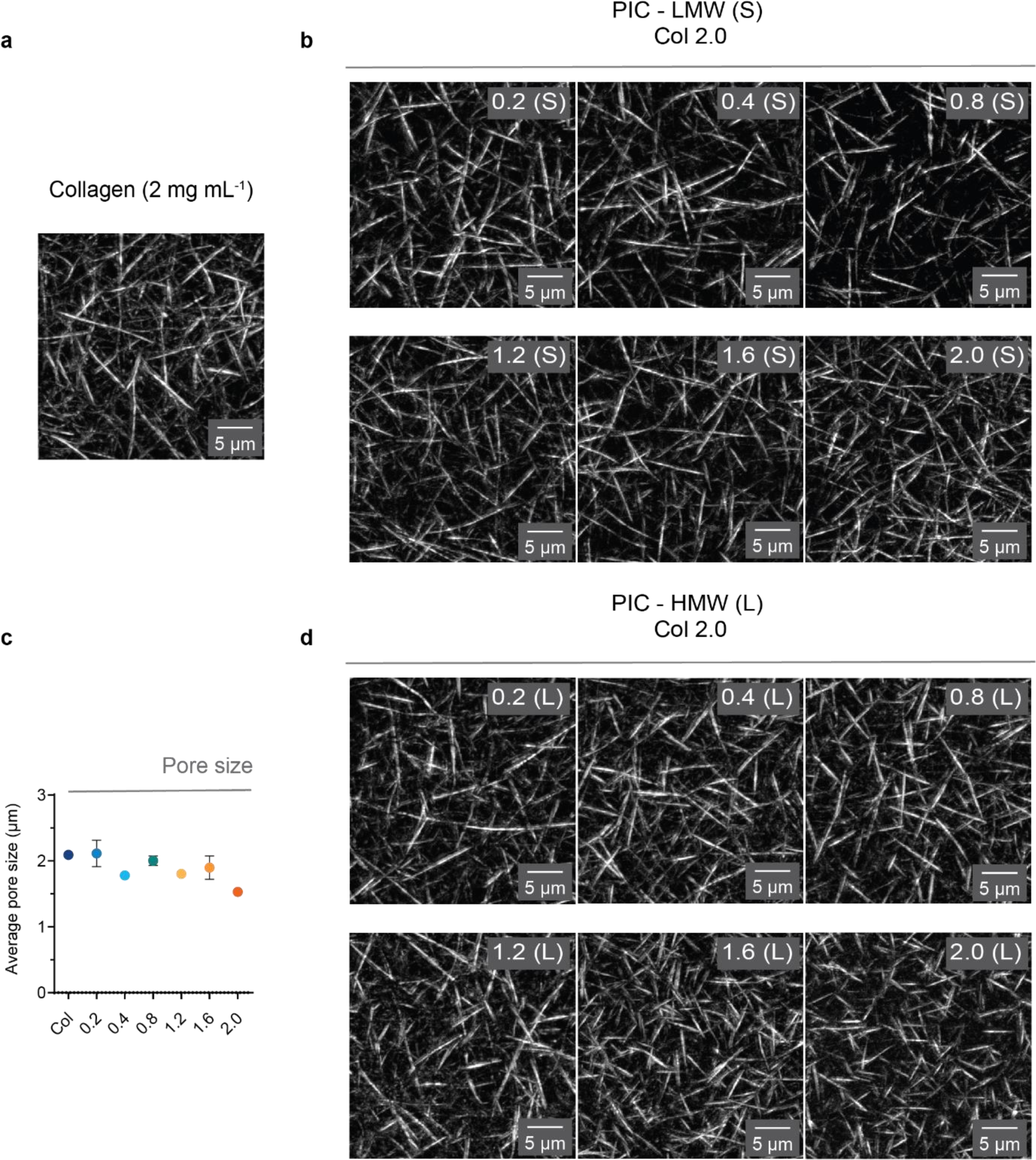
Pore size analysis by confocal laser scan microscopy. Z-projections of representative confocal reflection images of **a**, bovine atelo collagen (2 mg mL^−1^) and **b,** composites with increasing PIC concentrations (0.2–2.0 mg mL^−1^) using LMW PIC and **c-d,** using HMW PIC. Z-projections are 10 µm in depth. Scale bar (5 µm). **c,** Quantitative bubble analysis indicates no considerable change in the average pore size of bovine atelo collagen with increasing HMW PIC. Analysis shows data of three regions (z-projection of 50 µm depth each) taken subsequently within each hydrogel containing HMW PIC indicating homogeneity between the networks throughout the samples. Data represents an average of at least two independent measurements.

**Supplementary Fig 4.**
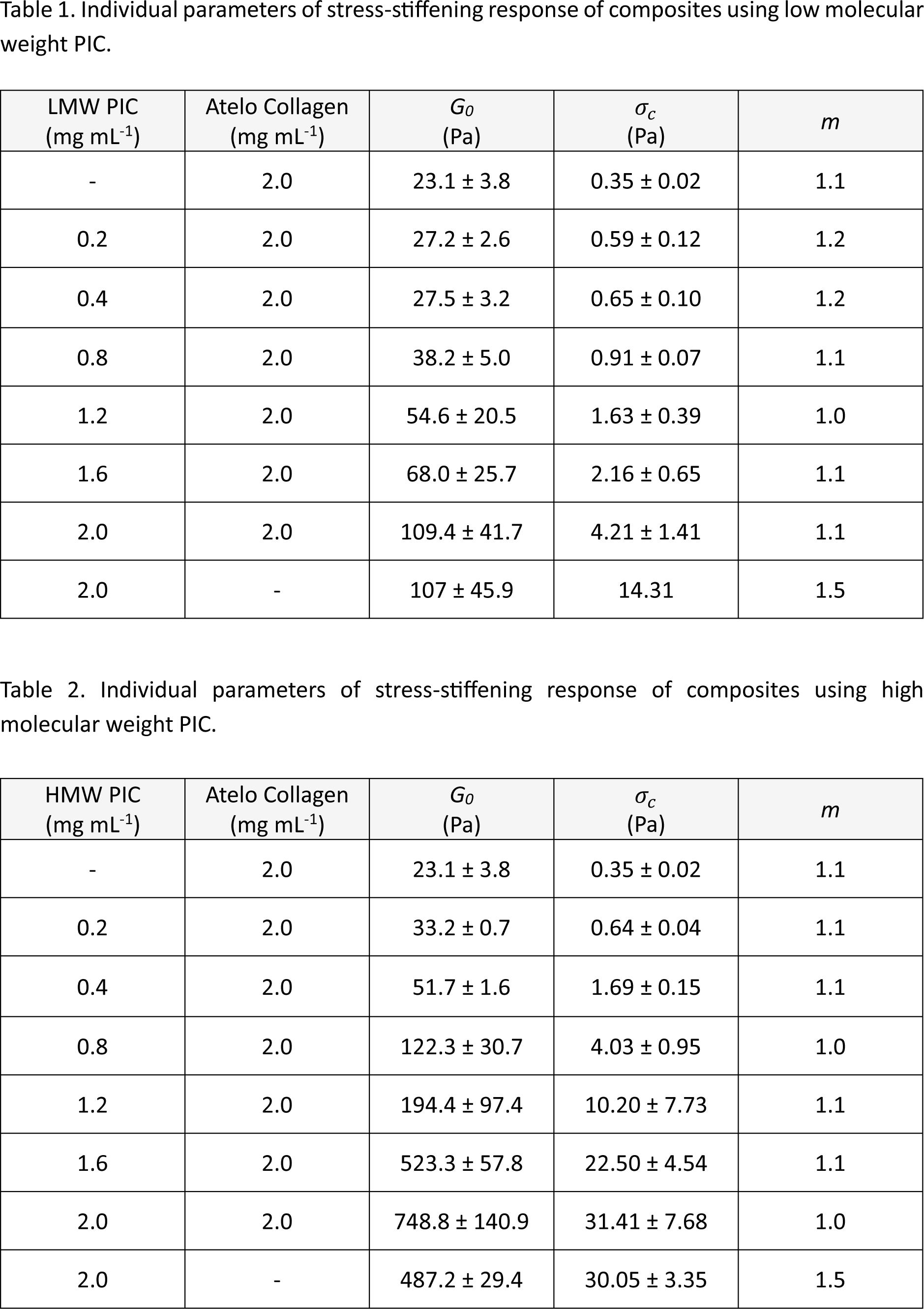
Extended Individual parameters of stress-stiffening response of collagen composites with LMW and HMW PIC.

**Supplementary Fig 5.**
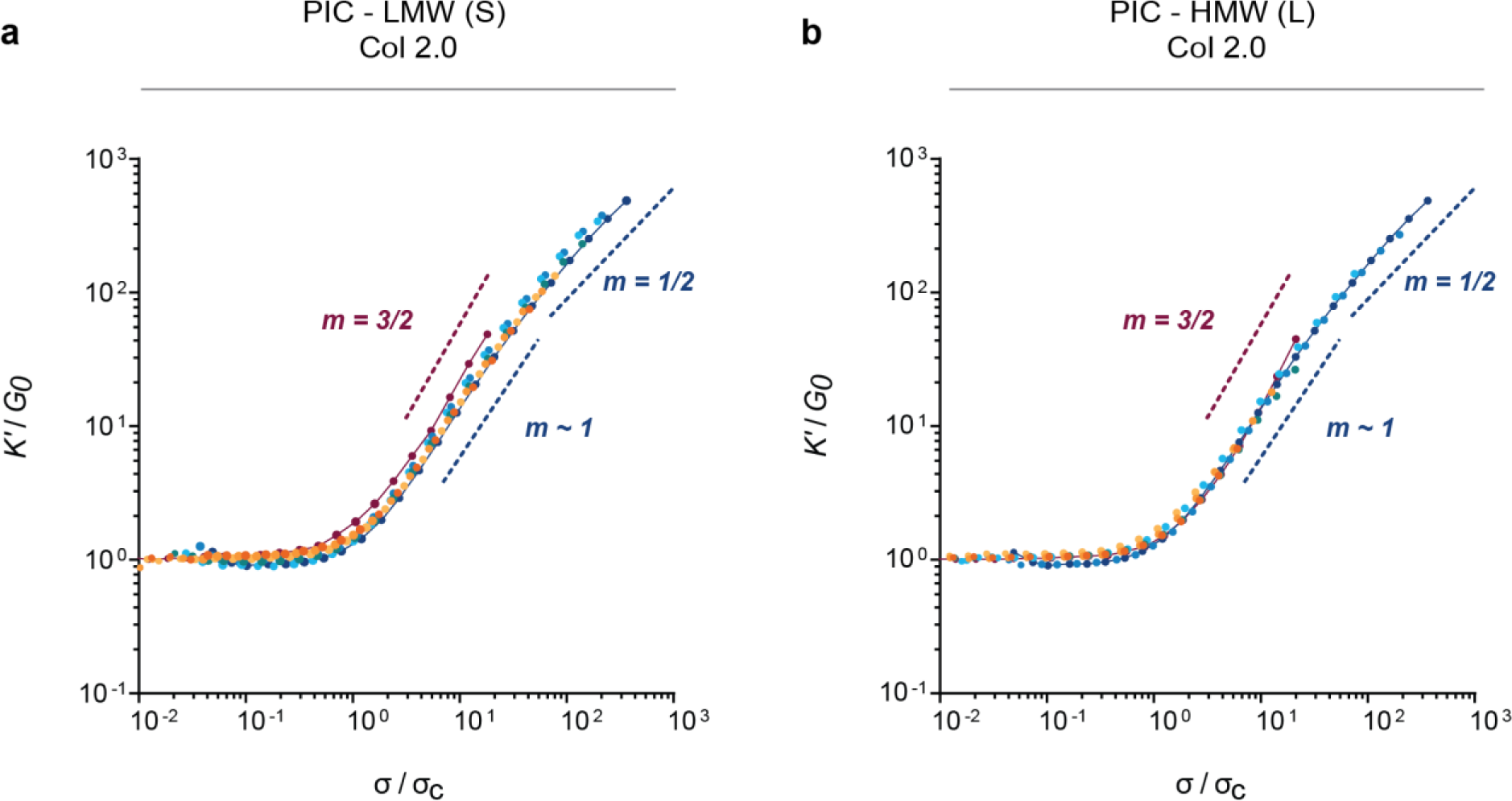
Fiber bending interactions dominate the non-linear response of PIC-Collagen composites. **a,b,** single master curves of PIC-Collagen composites were generated by scaling the differential modulus (*K’*) with the linear modulus (*G_0_*) and σ with σ_*c*_. The differential modulus (*K’*) of the PIC Collagen composites follows *K*’ ∝ σ^1^ characteristic of initial fibre bending of collagen network in the non-linear regime. Note that as the concentration of PIC increases, *K’* doesn’t follow an entropic stiffening where *K*′ ∝ σ^3/2^.

**Supplementary Fig 6.**
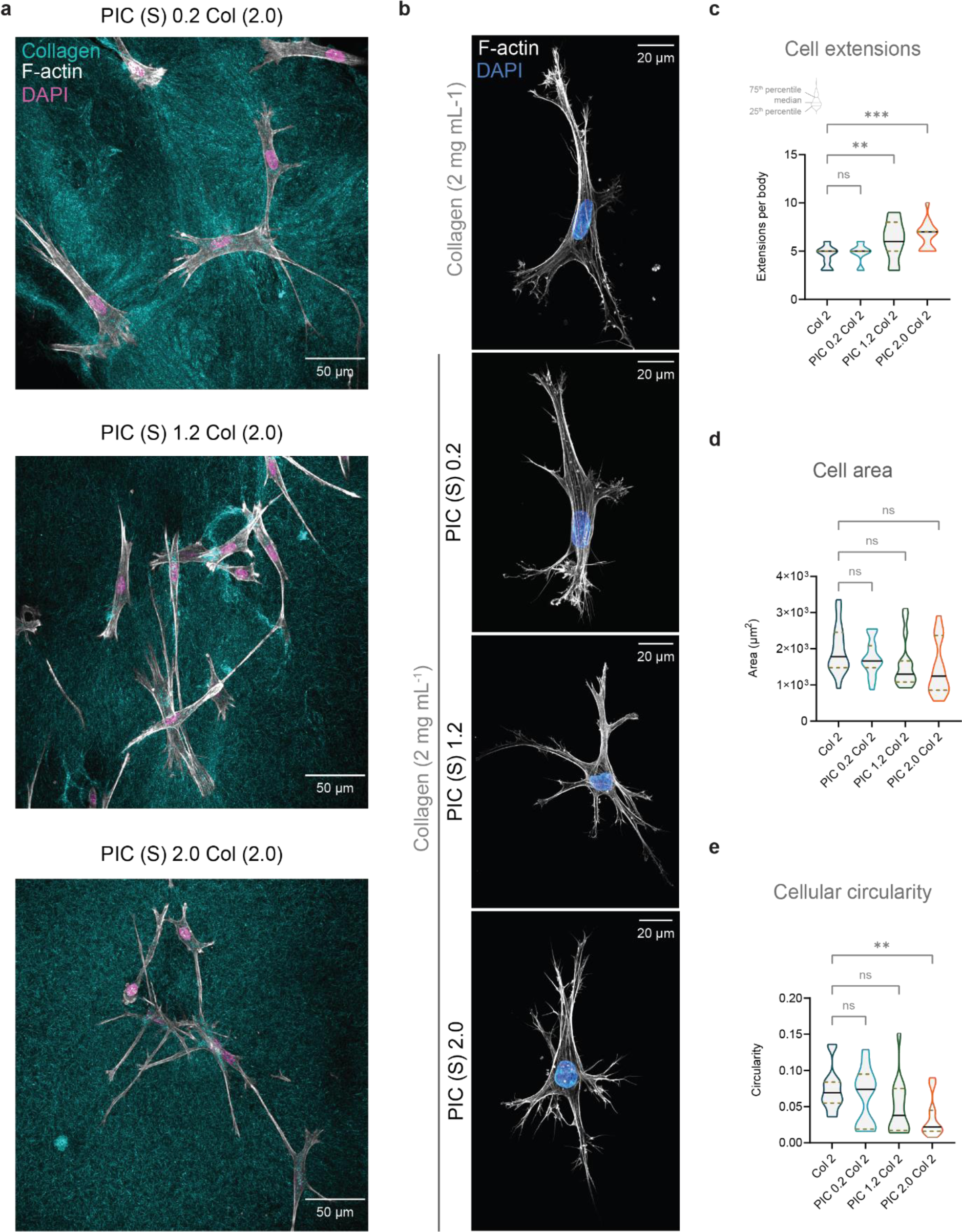
Collagen deformation in PIC Collagen composites and extended fibroblast characterisation. **a,** representative images of fibroblast morphology in cells embedded in PIC-Collagen composites (LMW) (0.2, 1.2, 2.0 mg mL^−1^). Samples were fixed 24 hours after embedding. The collagen network (cyan) images were acquired with confocal reflection mode. The cytoskeleton and nuclei were stained with Phalloidin 488 (white) and DAPI (magenta). Z projections are ∼50 µm depth. Samples exhibit a decrease in reflection intensity surrounding the leading edge of the cells with increasing PIC concentrations. **b,** extended representative images of fibroblasts and **c-e,** extended morphological quantification. Extended quantification includes **c,** number of extensions per body, **d,** cell body area (µm^2^), and **e,** cell circularity in each condition. Morphological features were obtained from a set of two independent measurements (n = 15 cells per condition). Violin plots (truncated, medium smooth) show median (black line) and quartiles (gold pattern lines).

**Supplementary Fig 7.**
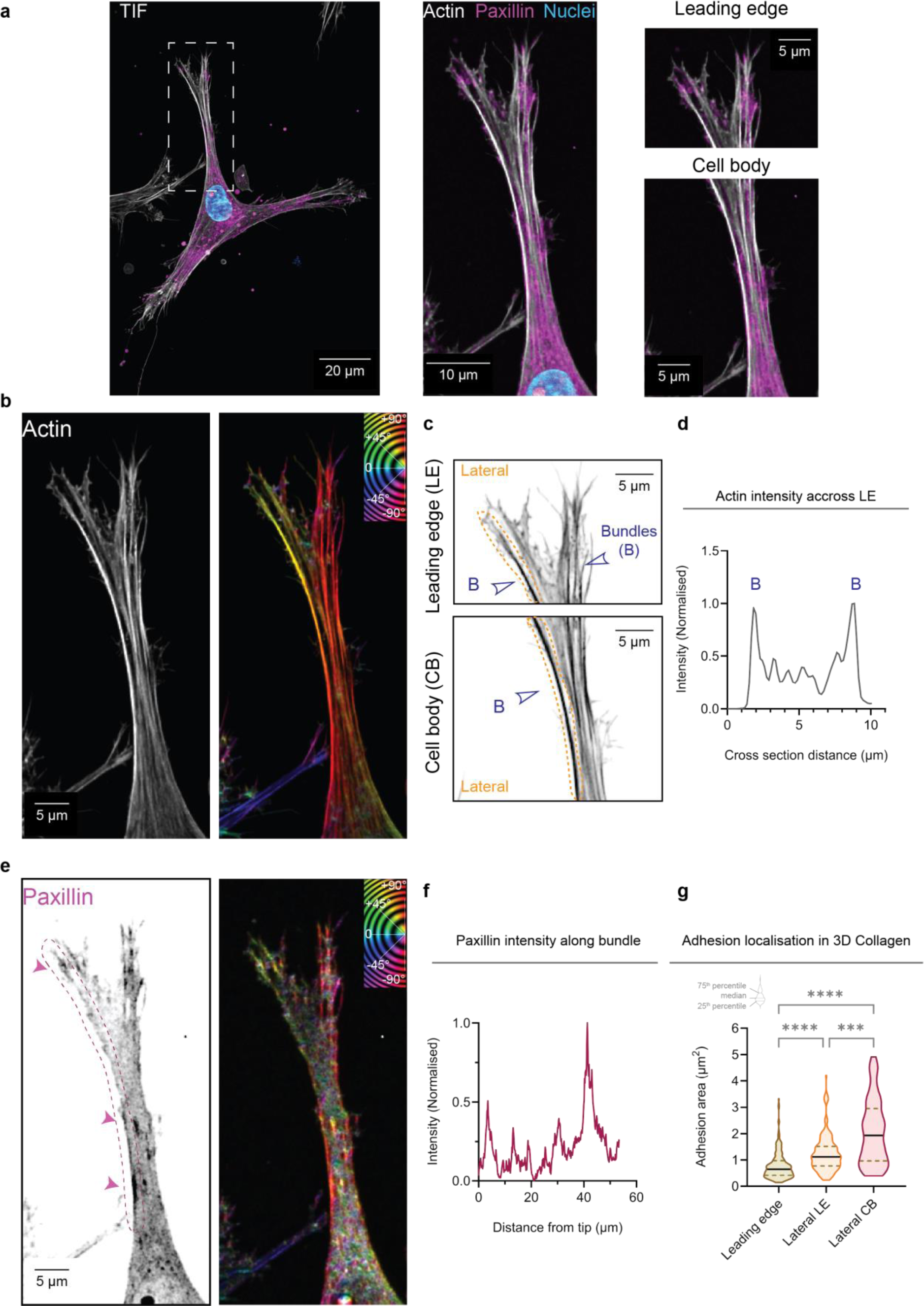
Spatial organisation of focal adhesions and the actin cytoskeleton in fibroblast cells embedded in 3D collagen hydrogels. **a,** confocal image of a fixed fibroblast expressing Paxillin–mCherry embedded in an atelo bovine collagen hydrogel (2 mg mL^−1^, 1X PBS, pH 7) after 24 hours. Actin and nuclei are stained with Phalloidin 488 and DAPI. Z projection ∼50 µm. Scale bar (20 µm). Inlet shows a zoomed region of a cell extension of interest (Scale bar (10 µm)) and the extension is classified into two subregions: leading edge and cell body. Scale bar (5 µm). **b,** organisation of the actin cytoskeleton at leading edge and cell body is depicted by the colour coded orientation (°, degrees) map of actin bundles and **c**, inverted confocal images. Both arrows and **d**, intensity profile (normalised) of a straight-line scan drawn across the body indicate an accumulation of F-actin localised at the lateral sides of both leading edge and cell body. Scale bar (5 µm). **e,** localisation of paxillin at both leading edge and lateral sides of leading edge/cell body shown by colour coded orientation of adhesions (paxillin) (°, degrees) and inverted confocal images. Note the matching colour coding of the adhesions and actin structures indicating an alignment and orientation of adhesions relative to the bundles. **f**, intensity profile (normalised) of a line scan drawn from tip to cell body (∼60 µm) following the direction of the actin bundle indicates different morphological phases of adhesions along the cell extension. **g**, adhesions at leading edge are smaller compared to adhesions at the lateral regions of the leading edge and cell body. Adhesions were manually quantified (Leading edge (n= 213), Lateral leading edge (n = 168), Lateral cell body (n = 69)) from a set of two independent measurements (30 cell extensions from 14 cells). Violin plots (truncated, medium smooth) show median (black line) and quartiles (gold pattern lines).

**Supplementary Fig 8.**
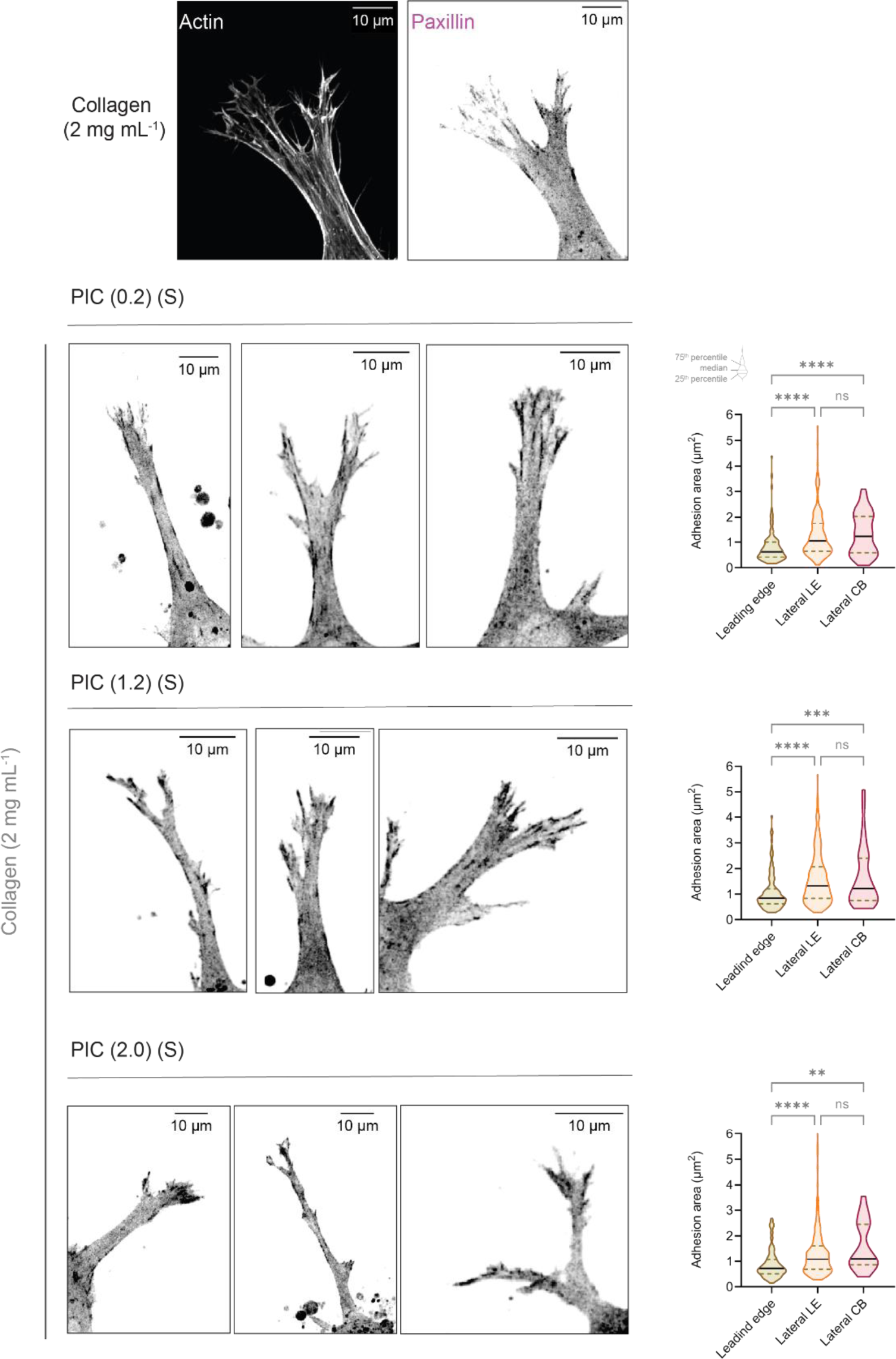
Extended data of cell-matrix interactions in PIC Collagen composites. Representative confocal images showing the localisation of paxillin at both leading edge and lateral sides of leading edge/cell body. In comparison to the localisation in pure collagen hydrogels, adhesions at the leading edge are smaller compared to adhesions at the lateral regions of the leading edge and cell body. Violin plots (truncated, medium smooth) show median (black line) and quartiles (gold pattern lines).

## References

1 Magnusson, S. P., Heinemeier, K. M. & Kjaer, M. Collagen Homeostasis and Metabolism. Metabolic Influences on Risk for Tendon Disorders, 11–25, doi:10.1007/978-3-319-33943-6_2.

2 Varga, F. et al. Biomechanical characterization of cartilages by a novel approach of blunt impact testing. Physiol Res 56, S61–S68, doi:10.33549/physiolres.931303 (2007).

3 Yang, W. et al. On the tear resistance of skin. Nat Commun 6, 6649–6649, doi:10.1038/ncomms7649 (2015).

4 Andrikakou, P., Vickraman, K. & Arora, H. On the behaviour of lung tissue under tension and compression. Sci Rep 6, 36642–36642, doi:10.1038/srep36642 (2016).

5 Shin, J.-W. et al. Molecular mechanisms of dermal aging and antiaging approaches. Int J Mol Sci 20, 2126, doi:10.3390/ijms20092126 (2019).

6 Jansen, K. A. et al. The Role of Network Architecture in Collagen Mechanics. Biophysical journal 114, 2665–2678 (2018).

7 Kjær, M. et al. Extracellular matrix adaptation of tendon and skeletal muscle to exercise. J Anat 208, 445–450, doi:10.1111/j.1469-7580.2006.00549.x (2006).

8 Miller, B. F. et al. Coordinated collagen and muscle protein synthesis in human patella tendon and quadriceps muscle after exercise. J Physiol 567, 1021–1033, doi:10.1113/jphysiol.2005.093690 (2005).

9 Saini, K., Cho, S., Dooling, L. J. & Discher, D. E. Tension in fibrils suppresses their enzymatic degradation – A molecular mechanism for ‘use it or lose it’. Matrix Biol 85-86, 34–46, doi:10.1016/j.matbio.2019.06.001 (2020).

10 van Helvert, S. & Friedl, P. Strain Stiffening of Fibrillar Collagen during Individual and Collective Cell Migration Identified by AFM Nanoindentation. ACS Appl. Mater. Interfaces 8, 21946–21955, doi:10.1021/acsami.6b01755 (2016).

11 Doyle, A. D., Sykora, D. J., Pacheco, G. G., Kutys, M. L. & Yamada, K. M. 3D mesenchymal cell migration is driven by anterior cellular contraction that generates an extracellular matrix prestrain. Developmental Cell 56, 826-+, doi:10.1016/j.devcel.2021.02.017 (2021).

12 Winer, J. P., Oake, S. & Janmey, P. A. Non-linear elasticity of extracellular matrices enables contractile cells to communicate local position and orientation. PLoS One 4, e6382–e6382, doi:10.1371/journal.pone.0006382 (2009).

13 Wang, H., Abhilash, A. S., Chen, Christopher S., Wells, Rebecca G. & Shenoy, Vivek B. Long-Range Force Transmission in Fibrous Matrices Enabled by Tension-Driven Alignment of Fibers. Biophys J 107, 2592–2603, doi:10.1016/j.bpj.2014.09.044 (2014).

14 Pakshir, P. et al. Dynamic fibroblast contractions attract remote macrophages in fibrillar collagen matrix.Nat Commun 10, 1850–1850, doi:10.1038/s41467-019-09709-6 (2019).

15 Wynn, T. A. Cellular and molecular mechanisms of fibrosis. J. Pathol 214, 199–210, doi:10.1002/path.2277 (2008).

16 Winkler, J., Abisoye-Ogunniyan, A., Metcalf, K. J. & Werb, Z. Concepts of extracellular matrix remodelling in tumour progression and metastasis. Nature communications 11, 5120–5120, doi:10.1038/s41467-020-18794-x (2020).

17 Potekaev, N. N. et al. The role of extracellular matrix in skin wound healing. J Clin Med 10, 5947, doi:10.3390/jcm10245947 (2021).

18 Kai, F., Drain, A. P. & Weaver, V. M. The Extracellular Matrix Modulates the Metastatic Journey. Dev Cell 49, 332–346, doi:10.1016/j.devcel.2019.03.026 (2019).

19 Schreml, S. M. D., Szeimies, R.-M. M. D. P., Prantl, L. M. D. P., Landthaler, M. M. D. P. & Babilas, P. M. D. P. Wound healing in the 21st century. J Am Acad Dermatol 63, 866–881, doi:10.1016/j.jaad.2009.10.048 (2010).

20 Xue, M. & Jackson, C. J. Extracellular Matrix Reorganization During Wound Healing and Its Impact on Abnormal Scarring. Adv Wound Care (New Rochelle) 4, 119–136, doi:10.1089/wound.2013.0485 (2015).

21 Yamauchi, M., Barker, T. H., Gibbons, D. L. & Kurie, J. M. The fibrotic tumor stroma. J Clin Invest 128, 16–25, doi:10.1172/JCI93554 (2018).

22 Laurens, N., Koolwijk, P. & De Maat, M. P. M. Fibrin structure and wound healing. J Thromb Haemost 4, 932–939, doi:10.1111/j.1538-7836.2006.01861.x (2006).

23 Voutouri, C. & Stylianopoulos, T. Accumulation of mechanical forces in tumors is related to hyaluronan content and tissue stiffness. PLoS One 13, e0193801–e0193801, doi:10.1371/journal.pone.0193801 (2018).

24 Tahkola, K. et al. Stromal hyaluronan accumulation is associated with low immune response and poor prognosis in pancreatic cancer. Sci Rep 11, 12216–12216, doi:10.1038/s41598-021-91796-x (2021).

25 Lai, V. K., Lake, S. P., Frey, C. R., Tranquillo, R. T. & Barocas, V. H. Mechanical behavior of collagen-fibrin co-gels reflects transition from series to parallel interactions with increasing collagen content. Journal of Biomechanical Engineering 134, doi:10.1115/1.4005544 (2012).

26 Lai, V. K. et al. Swelling of Collagen-Hyaluronic Acid Co-Gels: An In Vitro Residual Stress Model. Ann Biomed Eng 44, 2984–2993, doi:10.1007/s10439-016-1636-0 (2016).

27 Kim, O. V. et al. Compression-induced structural and mechanical changes of fibrin-collagen composites. Matrix Biol 60-61, 141–156, doi:10.1016/j.matbio.2016.10.007 (2017).

28 Nedrelow, D. S., Bankwala, D., Hyypio, J. D., Lai, V. K. & Barocas, V. H. Mechanics of a two-fiber model with one nested fiber network, as applied to the collagen-fibrin system. Acta Biomaterialia 72, 306–315, doi:10.1016/j.actbio.2018.03.053 (2018).

29 Burla, F., Tauber, J., Dussi, S. & Koenderink, G. Stress management in composite biopolymer networks. Nature Physics 15, 549–553 (2019).

30 Chen, X. et al. Glycosaminoglycans modulate long-range mechanical communication between cells in collagen networks. Proc Natl Acad Sci U S A 119, e2116718119–e2116718119, doi:10.1073/pnas.2116718119 (2022).

31 Kouwer, P. H. J. et al. Responsive biomimetic networks from polyisocyanopeptide hydrogels. Nature 493, 651, doi:10.1038/nature11839 (2013).

32 Jaspers, M. et al. Ultra-responsive soft matter from strain-stiffening hydrogels. Nat Commun 5, 5808–5808 (2014).

33 Das, R. J., Gocheva, V., Hammink, R., Zuani, O. F. & Rowan, A. E. Stress-stiffening-mediated stem-cell commitment switch in soft responsive hydrogels. Nature Materials 15, doi:10.1038/nmat4483 (2015).

34 Magno, V. et al. Macromolecular crowding for tailoring tissue-derived fibrillated matrices. Acta biomaterialia 55, 109–119, doi:10.1016/j.actbio.2017.04.018 (2017).

35 Dewavrin, J.-Y., Hamzavi, N., Shim, V. P. W. & Raghunath, M. Tuning the architecture of three-dimensional collagen hydrogels by physiological macromolecular crowding. Acta biomaterialia 10, 4351–4359 (2014).

36 Dewavrin, J.-Y. et al. Synergistic rate boosting of collagen fibrillogenesis in heterogeneous mixtures of crowding agents. The Journal of Physical Chemistry B 119, 4350–4358 (2015).

37 Ng, W. L., Goh, M. H., Yeong, W. Y. & Naing, M. W. Applying macromolecular crowding to 3D bioprinting: fabrication of 3D hierarchical porous collagen-based hydrogel constructs. Biomater. Sci. 6, 562–574 (2018).

38 Ranamukhaarachchi, S. et al. Macromolecular crowding tunes 3D collagen architecture and cell morphogenesis. Biomaterials science 7, 618–633 (2019).

39 Vandaele, J. et al. Structural characterization of fibrous synthetic hydrogels using fluorescence microscopy. Soft Matter 16, 4210–4219, doi:10.1039/c9sm01828j (2020).

40 Licup, A. J. et al. Stress controls the mechanics of collagen networks. Proceedings of the National Academy of Sciences 112, 9573–9578 (2015).

41 Sharma, A. et al. Strain-controlled criticality governs the nonlinear mechanics of fibre networks. Nature Physics 12, 584–587 (2016).

42 Ju, R. J., et al. Compression-dependent microtubule reinforcement comprises a mechanostat which enables cells to navigate confined environments. bioRxiv, 2022.2002.2008.479516, doi:10.1101/2022.02.08.479516 (2023).

43 Friedl, P., Wolf, K. & Lammerding, J. Nuclear mechanics during cell migration. Curr Opin Cell Biol 23, 55–64, doi:10.1016/j.ceb.2010.10.015 (2011).

44 Wolf, K. et al. Physical limits of cell migration: Control by ECM space and nuclear deformation and tuning by proteolysis and traction force. The Journal of cell biology 201, 1069–1084, doi:10.1083/jcb.201210152 (2013).

45 Davidson, M. W. et al. Nanoscale architecture of integrin-based cell adhesions. Nature 468, 580–584, doi:10.1038/nature09621 (2010).

46 Case, L. B. & Waterman, C. M. Integration of actin dynamics and cell adhesion by a three-dimensional, mechanosensitive molecular clutch. Nature Cell Biology 17, 955–963, doi:10.1038/ncb3191 (2015).

47 Zaidel-Bar, R., Ballestrem, C., Kam, Z. & Geiger, B. Early molecular events in the assembly of matrix adhesions at the leading edge of migrating cells. J Cell Sci 116, 4605–4613, doi:10.1242/jcs.00792 (2003).

48 Choi, C. K. et al. Actin and α-actinin orchestrate the assembly and maturation of nascent adhesions in a myosin II motor-independent manner. Nat Cell Biol 10, 1039–1050, doi:10.1038/ncb1763 (2008).

49 Gardel, M. L. et al. Traction Stress in Focal Adhesions Correlates Biphasically with Actin Retrograde Flow Speed. J Cell Biol 183, 999–1005, doi:10.1083/jcb.200810060 (2008).

50 Thievessen, I. et al. Vinculin-actin interaction couples actin retrograde flow to focal adhesions, but is dispensable for focal adhesion growth. J Cell Biol 202, 163–177, doi:10.1083/jcb.201303129 (2013).

51 Swaminathan, V. et al. Actin retrograde flow actively aligns and orients ligand-engaged integrins in focal adhesions. Proc Natl Acad Sci U S A 114, 10648–10653, doi:10.1073/pnas.1701136114 (2017).

52 Laukaitis, C. M., Webb, D. J., Donais, K. & Horwitz, A. F. Differential Dynamics of α5 Integrin, Paxillin, and α-Actinin during Formation and Disassembly of Adhesions in Migrating Cells. The Journal of cell biology 153, 1427–1440, doi:10.1083/jcb.153.7.1427 (2001).

53 Wiseman, P. W. et al. Spatial mapping of integrin interactions and dynamics during cell migration by image correlation microscopy. J Cell Sci 117, 5521–5534, doi:10.1242/jcs.01416 (2004).

54 Stehbens, S. J. & Wittmann, T. Analysis of focal adhesion turnover: A quantitative live-cell imaging example. Quantitative Imaging in Cell Biology 123, 335–346, doi:10.1016/B978-0-12-420138-5.00018-5 (2014).

55 Owen, L. M. et al. A cytoskeletal clutch mediates cellular force transmission in a soft, three-dimensional extracellular matrix. Molecular Biology of the Cell 28, 1959–1974, doi:10.1091/mbc.E17-02-0102 (2017).

56 Elosegui-Artola, A. et al. Mechanical regulation of a molecular clutch defines force transmission and transduction in response to matrix rigidity. Nat Cell Biol 18, 540–548, doi:10.1038/ncb3336 (2016).

57 Pankova, K., Rosel, D., Novotny, M. & Brabek, J. The molecular mechanisms of transition between mesenchymal and amoeboid invasiveness in tumor cells. Cell Mol Life Sci 67, 63–71, doi:10.1007/s00018-009-0132-1 (2010).

58 Taddei, M. L. et al. Mesenchymal to amoeboid transition is associated with stem-like features of melanoma cells. Cell Commun Signal 12, 24–24, doi:10.1186/1478-811X-12-24 (2014).

59 Holle, A. W. et al. Cancer Cells Invade Confined Microchannels via a Self-Directed Mesenchymal-to-Amoeboid Transition. Nano Lett 19, 2280–2290, doi:10.1021/acs.nanolett.8b04720 (2019).

60 Ju, R. J., Stehbens, S. J. & Haass, N. K. The Role of Melanoma Cell-Stroma Interaction in Cell Motility, Invasion, and Metastasis. Frontiers in Medicine 5, doi:10.3389/fmed.2018.00307 (2018).

61 Wisdom, K. M. et al. Matrix mechanical plasticity regulates cancer cell migration through confining microenvironments. Nat Commun 9, 4144–4113, doi:10.1038/s41467-018-06641-z (2018).

62 Infante, E. et al. LINC complex-Lis1 interplay controls MT1-MMP matrix digest-on-demand response for confined tumor cell migration. Nat Commun 9, 2443–2413, doi:10.1038/s41467-018-04865-7 (2018).

63 Howe, A. K., Baldor, L. C., Hogan, B. P. & Taylor, S. S. Spatial Regulation of the cAMP-Dependent Protein Kinase during Chemotactic Cell Migration. Proc Natl Acad Sci U S A 102, 14320–14325, doi:10.1073/pnas.0507072102 (2005).

64 Van Haastert, P. J. M. How cells use pseudopods for persistent movement and navigation. Sci Signal 4, pe6–pe6, doi:10.1126/scisignal.2001708 (2011).

65 Fritz-Laylin, L. K. et al. Actin-Based protrusions of migrating neutrophils are intrinsically lamellar and facilitate direction changes. Elife 6, doi:10.7554/eLife.26990 (2017).

66 Gierke, S. & Wittmann, T. EB1-Recruited Microtubule +TIP Complexes Coordinate Protrusion Dynamics during 3D Epithelial Remodeling. Curr Biol 22, 753–762, doi:10.1016/j.cub.2012.02.069 (2012).

67 Doyle, A. D., Carvajal, N., Jin, A., Matsumoto, K. & Yamada, K. M. Local 3D matrix microenvironment regulates cell migration through spatiotemporal dynamics of contractility-dependent adhesions. Nature Communications 6, doi:10.1038/ncomms9720 (2015).

68 Elosegui-Artola, A., Trepat, X. & Roca-Cusachs, P. Control of Mechanotransduction by Molecular Clutch Dynamics. Trends Cell Biol 28, 356–367, doi:10.1016/j.tcb.2018.01.008 (2018).

69 Sapoznik, E. et al. A versatile oblique plane microscope for large-scale and high-resolution imaging of subcellular dynamics. Elife 9, 1–39, doi:10.7554/eLife.57681 (2020).

70 Dutta, S. K., Mbi, A., Arevalo, R. C. & Blair, D. L. Development of a confocal rheometer for soft and biological materials. Rev Sci Instrum 84, 063702, doi:10.1063/1.4810015 (2013).

71 Tran-Ba, K.-H., Lee, D. J., Zhu, J., Paeng, K. & Kaufman, L. J. Confocal Rheology Probes the Structure and Mechanics of Collagen through the Sol-Gel Transition. Biophysical journal 113, 1882–1892 (2017).

72 Molteni, M., Magatti, D., Cardinali, B., Rocco, M. & Ferri, F. Fast two-dimensional bubble analysis of biopolymer filamentous networks pore size from confocal microscopy thin data stacks. Biophysical journal 104, 1160–1169 (2013).

73 Münster, S. & Fabry, B. A simplified implementation of the bubble analysis of biopolymer network pores. Biophysical journal 104, 2774–2775 (2013).

74 Gilbert, E. P., Schulz, J. C. & Noakes, T. J. ’Quokka’ - the small-angle neutron scattering instrument at OPAL. Physica B 385-86, 1180–1182, doi:10.1016/j.physb.2006.05.385 (2006).

75 Rehm, C. et al. Design and performance of the variable-wavelength Bonse–Hart ultra-small-angle neutron scattering diffractometer KOOKABURRA at ANSTO. J Appl Crystallogr 51, 1–8, doi:10.1107/S1600576717016879 (2018).

76 Kline, S. R. Reduction and analysis of SANS and USANS data using IGOR Pro. J Appl Crystallogr 39, 895–900, doi:10.1107/S0021889806035059 (2006).

77 Wood, K. et al. QUOKKA, the pinhole small-angle neutron scattering instrument at the OPAL Research Reactor, Australia: design, performance, operation and scientific highlights. J Appl Crystallogr 51, 294–314, doi:10.1107/S1600576718002534 (2018).

78 Gavrilov, M., Gilbert, E. P., Rowan, A. E., Lauko, J. & Yakubov, G. E. Structural Insights into the Mechanism of Heat-Set Gel Formation of Polyisocyanopeptide Polymers. Macromolecular rapid communications. 41, 2000304-n/a, doi:10.1002/marc.202000304 (2020).

79 Stehbens, S. J. et al. CLASPs link focal-adhesion-associated microtubule capture to localized exocytosis and adhesion site turnover. Nature Cell Biology 16, 558-+, doi:10.1038/ncb2975 (2014).

